# Morphogenesis and cell ordering in confined bacterial biofilms

**DOI:** 10.1101/2021.02.17.431682

**Authors:** Qiuting Zhang, Jian Li, Japinder Nijjer, Haoran Lu, Mrityunjay Kothari, Ricard Alert, Tal Cohen, Jing Yan

## Abstract

Biofilms are aggregates of bacterial cells surrounded by an extracellular matrix. Much progress has been made in studying biofilm growth on solid substrates; however, little is known about the biophysical mechanisms underlying biofilm development in three-dimensional confined environments, in which the biofilm-dwelling cells must push against and even damage the surrounding environment to proliferate. Here, combining single-cell imaging, mutagenesis, and rheological measurement, we reveal the key morphogenesis steps of *Vibrio cholerae* biofilms embedded in hydrogels as they grow by four orders of magnitude from their initial size. We show that the morphodynamics and cell ordering in embedded biofilms are fundamentally different from those of biofilms on flat surfaces. Treating embedded biofilms as inclusions growing in an elastic medium, we quantitatively show that the stiffness contrast between the biofilm and its environment determines biofilm morphology and internal architecture, selecting between spherical biofilms with no cell ordering and oblate ellipsoidal biofilms with high cell ordering. When embedded in stiff gels, cells self-organize into a bipolar structure that resembles the molecular ordering in nematic liquid crystal droplets. *In vitro* biomechanical analysis shows that cell ordering arises from stress transmission across the biofilm-environment interface, mediated by specific matrix components. Our imaging technique and theoretical approach are generalizable to other biofilm-forming species, and potentially to biofilms embedded in mucus or host tissues as during infection. Our results open an avenue to understand how confined cell communities grow by means of a compromise between their inherent developmental program and the mechanical constraints imposed by the environment.

**Significance Statement:** Biofilms are microbial cities in which bacterial cells reside in a polymeric matrix. They are commonly found inside soft confining environments such as food matrices and host tissues, against which bacteria must push to proliferate. Here, by combining single-cell live imaging and mechanical characterization, we show that the confining environment determines the dynamics of biofilm shape and internal structure. This self-organized evolution of biofilm architecture is caused by force transmission between the environment and the biofilm, mediated by the extracellular matrix secreted by the cells. Our findings lead to new ways to understand how bacterial communities develop under mechanical constraints, and potentially to new strategies for preventing and controlling biofilm growth in three-dimensional environments.

## Introduction

The growth of living organisms is critically influenced by the external environment. One form of such environmental influence is the transmission of mechanical stress, which can instruct morphogenesis in systems ranging from stem cells to tissues to the entire organisms (1, 2). In the prokaryotic domain, bacteria commonly live in complex communities encased by an extracellular matrix (3), known as biofilms (4). Biofilm formation is a morphogenetic process whereby a single founder cell develops into a three-dimensional aggregate in which bacterial cells interact with each other and with the environment (4–8). Recent work has revealed biophysical mechanisms underlying biofilm morphogenesis on solid substrates, which is controlled by cell-substrate adhesion and the resulting shear stress (9–15). In addition to those living on surfaces, bacterial communities are also commonly found inside soft, structured environments, such as hydrogels. Examples include biofilms growing in mucus layers and host tissues during an infection or food contamination (16). Indeed, many common biofilm formers including *Pseudomonas aeruginosa* and *Vibrio cholerae* encounter biological hydrogels as their niche during infection (17, 18). Under these conditions, embedded biofilms must grow against three-dimensional (3D) confinement and potentially damage the surrounding environment — a process that is fundamentally different from how surface-attached biofilms expand against friction with the surface (10, 13, 15, 19). However, little is known about how biofilms develop under such 3D mechanical constraints, including how cells collectively organize in response to the confinement and how the confining environment, in turn, is modified by cell proliferation. This is in stark contrast to the accumulating knowledge on the growth dynamics of mammalian cell aggregates and tumors under confinement (20, 21).

In this study, we integrate single-cell live imaging, mutagenesis, *in vitro* mechanical testing, and numerical modeling to investigate how the 3D confinement determines the morphodynamics and cell ordering of an embedded biofilm. A model system is established by embedding *V. cholerae*, the causal agent of the cholera pandemic and a model biofilm former (22, 23), inside agarose gels (24). By using 3D visualization techniques with high spatiotemporal resolution, we reveal that embedded biofilms undergo a shape transition and a series of self-organization events that are distinct from those in surface-attached biofilms. We first show that the stiffness contrast between the biofilm and the confining hydrogels controls a transition between spherical and ellipsoidal biofilms. Furthermore, we discover that embedded biofilms display a core-shell structure with intricate ordering similar to nematic liquid crystal droplets (25). Finally, we demonstrate that Vibrio polysaccharide (VPS) and cell-to-surface adhesion proteins effectively transmit stress between the environment and the biofilm, giving rise to distinct cell ordering patterns in embedded biofilms.

## Results

### Single-cell imaging reveals distinct growth pattern of embedded *V. cholerae* biofilms

A convenient agarose-based assay is used to investigate the interplay between biofilm growth and mechanical confinement: individual cells are embedded at low density and allowed to grow into isolated biofilm clusters inside agarose gels (**Fig. 1A** and Fig. S1). We systemically vary the stiffness of the embedding gels from 100 Pa to 100 kPa by controlling the agarose concentration, a range that covers many soft biological hydrogels (1, 26). The biocompatibility and tunable mechanical properties render agarose gels a suitable embedding medium to grow biofilms without affecting the growth rate of the encapsulated cells (Fig. S2). We use the well-characterized rugose (Rg) *V. cholerae* strain locked in a high cyclic diguanylate level to focus on the biomechanical aspect of biofilm growth (9, 27). We start with a mutant lacking the cell-to-cell adhesion protein RbmA (23), as the surface-attached biofilm from this mutant is best understood (10, 11, 13). To quantitatively track the architecture of growing biofilms, we adopt an adaptive imaging technique to obtain microscopy data with high spatiotemporal resolution over the full course of biofilm growth (9, 11). We are able to visualize both the global morphology and the single-cell level architecture of an embedded biofilm containing up to 10^4^ cells (**Fig. 1, B and C**, Movie S1) with minimal phototoxicity and photobleaching. In parallel, we develop an adaptive local thresholding algorithm to segment individual cells in 3D confined biofilms (**Fig. 1D** and Fig. S3, Movie S2). This new algorithm overcomes the issue of image deterioration due to dense cellular packing and light scattering, both of which are significant for embedded biofilms.

**Fig.1.**
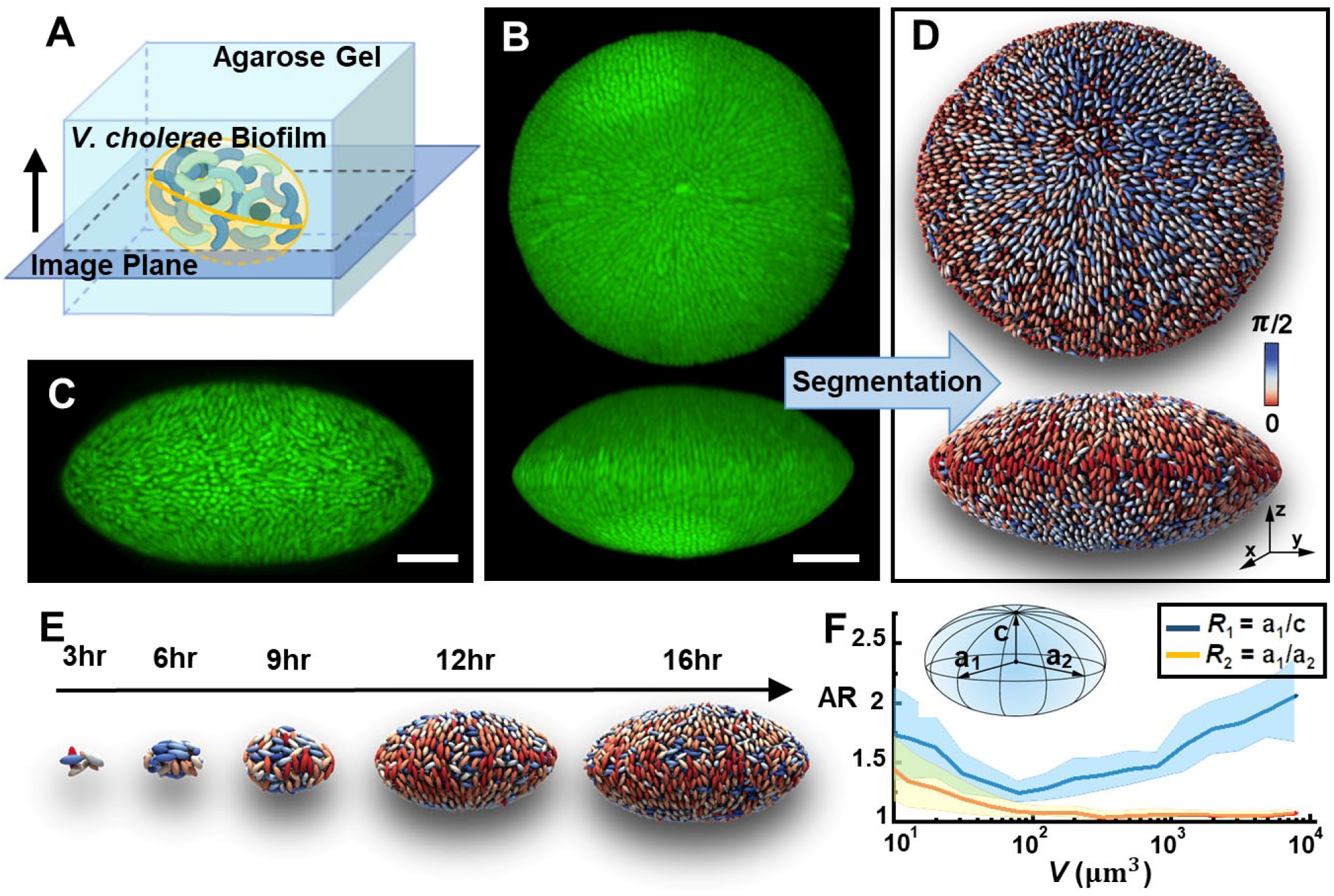
Single-cell imaging of embedded *V. cholerae* biofilms. (*A*) Schematic illustration of biofilm growth in agarose gels. (*B*) 3D view (top and side) and (*C*) cross-sectional image of a *V. cholerae* biofilm embedded in a 2% agarose gel. Cells constitutively express mNeonGreen. Scale bar: 10 μm. (*D*) Single-cell 3D reconstruction of the embedded biofilm shown in (*B*). Cells are color-coded according to their angles with respect to the *z* axis. (*E*) Time-lapse images of reconstructed biofilms. Images are rescaled differently at each time point for clarity. (*F*) Evolution of biofilm morphology inside 2% agarose gel characterized by two aspect ratios (ARs) as a function of biovolume *V*. Inset shows the definition of aspect ratios *R*_1_ = *a*_1_/*c* and *R*_2_ = *a*_1_/*a*_2_, in which *a*_1_, *a*_2_, and *c* (*a*_1_ > *a*_2_ > *c*) are the biofilm’s principal axes. Center lines correspond to means; widths of the shaded areas correspond to standard deviation (SDs, *n* = 10).

Both on global and single-cell levels, mature embedded biofilms possess a 3D organization distinct from that in biofilms growing on solid substrates. At the global level, the entire biofilm develops an anisotropic oblate shape. This is unexpected because previous results showed that free-floating biofilm clusters are roughly spherical (9). At the single-cell level, embedded biofilms display a precise pattern of nematic cell alignment: at the biofilm-gel interface, the rod-shaped *V. cholerae* cells align radially around the two poles of the biofilm. In the language of liquid crystals (LCs), the two poles correspond to +1 defects around which the local alignment axis rotates by 360° (28). The observed configuration is reminiscent of the bipolar organization of LC molecules on the surface of droplets, with the two +1 defects known as Boojum points (25).

Our imaging pipeline also reveals the key developmental stages of embedded biofilms. We follow the growth of well-separated biofilms from single founder cells to mature clusters at a time resolution of one cell division cycle (**Fig. 1E**, Fig. S4, and Movie S3). The reconstructed data at single-cell resolution show that, initially, small clusters are elongated due to the inherent shape anisotropy of the founder cell and the subsequent division along its long axis. At the early stage, embedded biofilms tend to grow from a prolate shape (*R*_1_>*R*_2_ >1, **Fig. 1F**) to a close-to-spherical shape (*R*_1_~*R*_2_~1). However, as the biofilm keeps growing, the spherical shape gradually transforms into an oblate shape (*R*_1_>*R*_2_ =1). Concomitantly, cells at the biofilm-gel interface develop the bipolar nematic order described above. The nonmonotonic biofilm shape evolution and the bipolar cell ordering pose a central question: *How does mechanical confinement impact biofilm growth and architecture both at global and local levels?*

### Biofilm-environment stiffness contrast determines biofilm morphology

We hypothesize that the anisotropic biofilm shapes result from the accumulation of growth-induced mechanical stresses under the constraints imposed by the environment. Therefore, we measure the evolution of the major aspect ratio *R*_1_ versus volume of biofilms subjected to different degrees of confinement by tuning agarose stiffness (**Fig. 2A**). The morphological evolution depends strongly on the degree of confinement: when the gel concentration is low (≤0.5%), the biofilm grows towards a spherical shape and stays spherical. In contrast, biofilms in stiffer hydrogels undergo a prolate-to-spherical-to-oblate transition and mature biofilms maintain an oblate shape.

**Fig.2.**
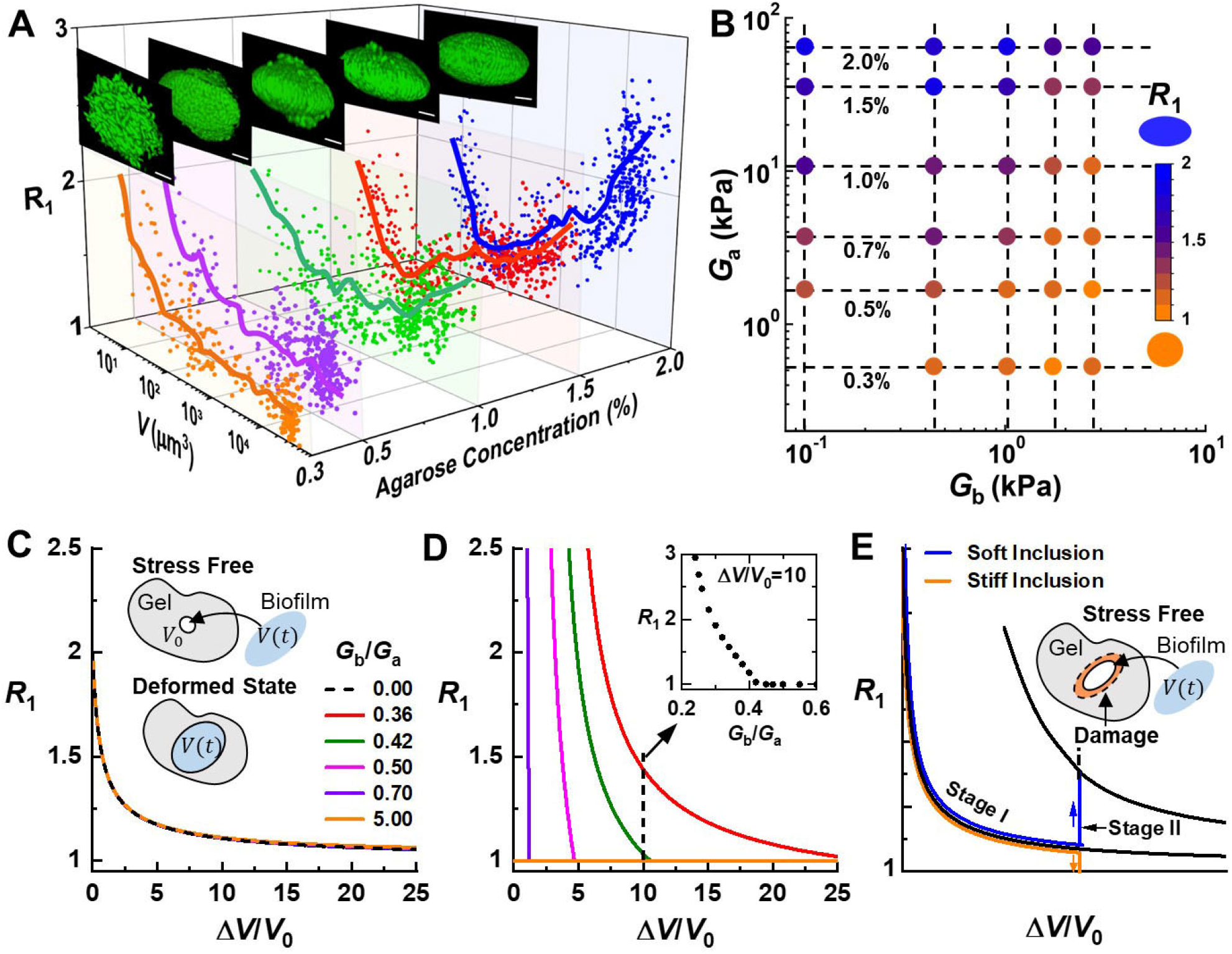
Stiffness contrast determines biofilm shape anisotropy. (*A*) Morphologies of biofilms growing in hydrogels solidified with different concentrations of agarose. Plotted are the biofilm aspect ratio *R*_1_ versus *V* obtained at different developmental stages. Solid lines are running means of the data points. Insets show images of representative biofilms for each condition. Scale bar: 5 μm. (*B*) Diagram showing biofilm shape as a function of biofilm stiffness *G*_b_ and agarose gel stiffness *G*_a_. Symbols are color-coded according to the average *R*_1_ of mature biofilms, ranging from 1 (spherical) to 2 (oblate). (*C-E*) Theoretical modeling examines *R*_1_ variation as a function of the volumetric expansion ratio (Δ*V/V*_0_) for different stiffness contrasts *G*_b_/*G*_a_ and at different stages of expansion. *V*0 corresponds to initial void volume and Δ*V* = *V* − *V*_0_. (*C*) Stage I – purely elastic deformation of the gel without any damage. Insets illustrate the modeling approach to determine the optimal biofilm shape. (*D*) Stage II – onset of damage in the gel environment. Inset shows that at a fixed volumetric ratio Δ*V/V*_0_, *R*_1_ decreases with increasing stiffness contrast *G*_b_/*G*_a_, qualitatively capturing the experimental observations in (*B*). (*E*) Schematic explanation of the shape evolution of embedded biofilms: In Stage I, the gel deforms elastically, and *R*_1_ evolves toward 1 for both soft and stiff inclusions (biofilms), similar to the expansion of a fluid cavity. After gel damage (indicated by the orange region) occurs in Stage II, *R*_1_ increases for soft inclusions but drops to 1 for stiff inclusions.

We envision that embedded biofilm growth bears conceptual similarity with the classical problem of elastic cavitation, i.e. the expansion of fluid cavity inside an elastic material (29–31). However, unlike liquid inclusions, a biofilm behaves as a viscoelastic and poroelastic solid (32); it resists deformations imposed by the confining environment to some extent (Fig. S5). Therefore, embedded biofilm growth can be conceptualized as a *solid inclusion* expanding inside another *solid*. The mechanical properties of the biofilm and the agarose gel can be characterized by their shear moduli, *G*_b_ and *G*_a_ (tables S1 and S2), respectively. We vary the stiffness of a biofilm by using different mutants lacking one or more extracellular matrix components (33). We measure the average aspect ratio of mature biofilms (~10^4^ μm^3^) for different stiffness contrasts between the biofilm and the embedding hydrogel, *G*_b_/*G*_a_ (**Fig. 2B**). The resulting diagram shows that the anisotropic biofilm morphology is favored when the environment is stiffer than the embedded biofilm (*G*_b_/*G*_a_ < 1, **Fig. 2B**).

### Continuum modeling of inclusion growth reveals that gel damage causes a transition in biofilm morphology

As the embedded biofilms grow by four orders of magnitude from their initial size, the surrounding materials experience tremendous deformation (30, 34) and mechanical stress beyond the elastic limit of the gel (Fig. S6 and Table S3). To explain the observed biofilm shape evolution, we develop a continuum model that couples biofilm growth with the nonlinear mechanical response of the environment (35). We model the biofilm as a solid inclusion constrained to fit into a void in an infinite elastic medium, with biofilm growth driving void expansion (**Fig. 2C, inset**, Fig. S7-9). If the biofilm were not subject to the confinement, it would develop a random organization resulting from cell proliferation in all directions (9). This indicates the absence of an inherent, programmed morphology in growing biofilms, in contrast to higher-order organisms. Hence, the key factor in determining biofilm shape is the accumulation of mechanical energy imposed by the external constraint. Specifically, we hypothesize that an embedded biofilm will evolve in a way to minimize the total mechanical energy in the system. This energy minimization yields an optimal shape of the confined biofilm at every given volume; this optimal shape is different from the stress-free shapes of both the void and the biofilm (Fig. S10).

Given that the surrounding material endures large strains, we propose two growth stages. In Stage I (**Fig. 2C**), the confining gel deforms elastically so that the stress-free shape of the void is preserved while the biofilm is allowed to vary its stress-free shape. Our calculations show that in this stage the optimal *R*_1_ quickly decreases to 1 regardless of the stiffness contrast *G*_b_/*G*_a_ – a situation similar to the classical problem of fluid cavity expansion. In Stage II, the strain generated by biofilm growth exceeds the elastic limit of the gel material in the region immediately surrounding the biofilm – damage thus occurs (34). Accordingly, the void can no longer maintain its original stress-free shape and we assume that the void’s stress-free shape now varies with that of the growing biofilm (**Fig. 2D**). Therefore, we predict that the biofilm adopts a new optimal shape at the onset of damage. This hypothesis is corroborated by our experiments showing that the volume at which the shape transition takes place scales with the yield strain of the gel (**Fig. 2A** and Fig. S11). It should be noted that the observed experimental transitions are expected to be gradual due to multiple relaxation processes in both the biofilm and the environment. Interestingly, this transition strongly depends on *G*_b_/*G*_a_ (**Fig. 2D, inset**). Combining the results of Stage I and II, we interpret the full evolution of biofilm shape (**Fig. 2E**). Starting from an elongated shape, an embedded biofilm initially evolves toward a spherical shape as if it were a fluid (Stage I); upon the onset of gel damage (Stage II), a biofilm stiffer than its environment will become even more spherical, while a biofilm softer than the environment will transition to an oblate shape.

### Bacterial cells self-organize as embedded biofilms grow

As shown in **Fig. 1**, cells at the biofilm-gel interface self-organize into an ordered pattern under confinement. To quantify cell ordering, we employ established tools in LCs (28). We measure cell orientation with respect to the biofilm-gel interface to define two orthogonal order parameters (Fig. S12): the shell order parameter *S*_s_ and the bipolar order parameter *S*_b_. *S*_s_ is defined from the angle between the cell director and the local surface normal, and it varies from 0 for a random configuration to 1 for cells lying parallel to the interface. To characterize the ordering *in* the plane locally tangential to the interface, we make an analogy between biofilm architecture and the arrangement of molecules on the surface of a LC droplet. In the latter case, Boojum points (+1 defects) are observed at two poles (25), from which molecules emanate radially and follow the lines of constant longitude (meridians) throughout the biofilm surface. Thus, we use the bipolar order parameter *S*_b_ to quantify how cells align with the local meridian.

We average *S*_s_ and *S*_b_ over cells in the two outmost layers to characterize global cell ordering near the interface. We follow the time evolution of 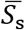 and 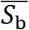 (**Fig. 3A** and Movie S4). As embedded biofilms develop, both 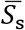 and 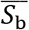 increase steadily with biofilm volume, indicating a gradual self-organization process. Interestingly, the increase in cell ordering is tightly coupled to the global morphological transition. Late in Stage I, accumulating pressure from biofilm expansion favors tangential alignment of cells at the biofilm-gel interface, leading to a fast increase in 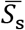. In contrast, 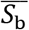 starts to increase only after 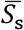 reaches a value ~0.5. We therefore conclude that shell ordering is a prerequisite for bipolar ordering. Bipolar ordering does not develop until the biofilm enters Stage II in which its oblate shape geometrically defines two poles.

**Fig.3.**
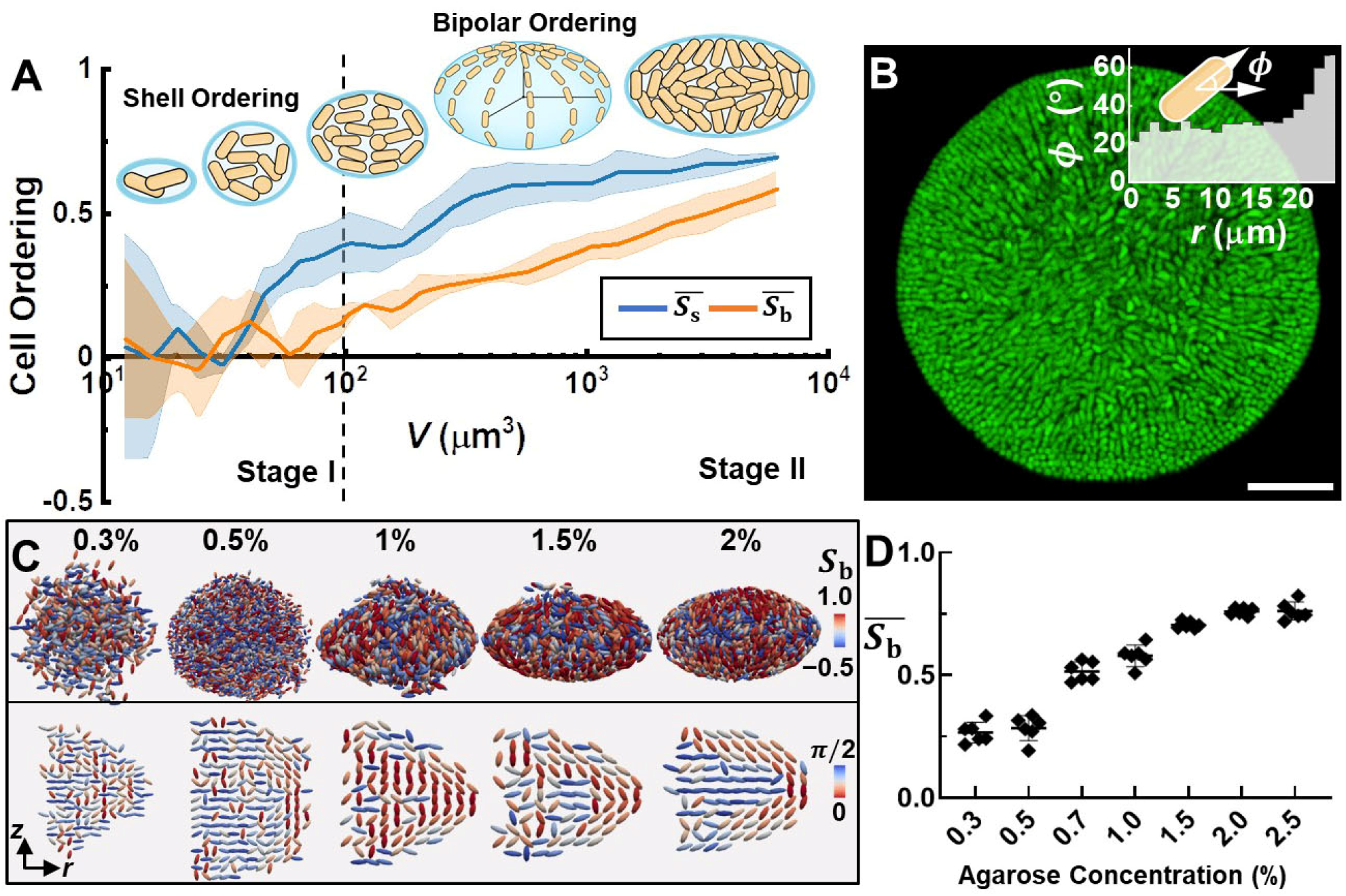
Cell ordering is promoted by biofilm-gel mechanical interactions. (*A*) Evolution of cell ordering with biovolume *V* for a biofilm in 2% gel. Cell ordering is quantified by the average shell ordering 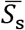 and average bipolar ordering 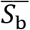 in the outmost cell layers. Center lines correspond to means; widths of the shaded areas correspond to SDs (*n* = 3). Inset: Schematic representation of the development of biofilm architecture. (*B*) Cross-sectional view of the principal plane of a mature biofilm. Inset: Average angle *ϕ* between a cell’s axis and the biofilm’s principal plane as a function of the distance from the center *r*. Scale bar: 10 μm. (*C*) Architecture of mature biofilms inside gels of designated agarose concentrations. Shown are the 3D reconstructed biofilms with each cell colored according to its bipolar order parameter *S*_b_ (*Top*) and average angle between the cell axis and the *z* axis (*Bottom*, see Fig. S12). (*D*) Average bipolar order parameter 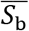 of mature biofilms grown in agarose gels with the designated concentrations. Error bars correspond to SDs (*n* = 7).

The interior of an embedded biofilm is much less ordered (**Fig. 3B**). However, we do observe a tendency for cells in the core to lie at an average angle of 20° with respect to the principal plane of the biofilm (**Fig. 3B, inset**). Interestingly, there is no preferred orientation within that plane, indicating that cells in the interior are shielded from the shell ordering at the interface (Fig. S13).

### Cell ordering is promoted by mechanical interactions across the biofilm-environment interface

We further investigate the dependence of cell ordering on gel stiffness (**Fig. 3C** and Fig. S14). The azimuthally averaged cell orientation (**Fig. 3C, *Bottom***) shows that both the bipolar organization of cells at the biofilm-gel interface and the planar arrangement at the biofilm core become more pronounced as the gel stiffness increases. Two factors underlie the disordered configuration for biofilms grown in low concentration gel (<1%). First, due to the large pore size of these gels (36), cells at the biofilm-gel interface are not necessarily tangential to the interface. Second, biofilms grown in low concentration gels experience weaker compression as evidenced by their lower cellular density (Fig. S2). Under these conditions, both 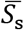 and 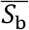 remain low. As the gel concentration increases, its stiffness increases and so does the mechanical resistance experienced by the embedded biofilm. Accordingly, biofilm cell density increases, and the biofilm shape becomes oblate. The concomitant increase in 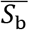 suggests that mechanical stress plays a critical role in cell ordering in confined biofilms (**Fig. 3D**).

### Extracellular matrix mediates cell ordering by transmitting stress from the environment

Previous studies on biofilms growing on rigid substrates have shown that cell ordering depends critically on cell-to-surface adhesion (9, 10), mediated by VPS and accessory proteins RbmC and Bap1 (37–39). We hypothesize that these matrix components also contribute to the cell ordering in embedded biofilms. To test this hypothesis, we generate mutants lacking one or more matrix components (**Fig. 4, A** and **B**) (9, 33). We observe that the colony from the Δ*vpsL* mutant unable to produce VPS maintains significant shell order (**Fig. 4A** and Fig. S14), suggesting that the tendency of cells to align parallel to the biofilm-gel interface is a generic result of confinement and independent of the biofilm matrix. However, Δ*vpsL* cells at the biofilm-gel interface fail to align in a bipolar order, and the biofilm core remains disordered. The Δ*rbmA*Δ*bap1*Δ*rbmC* (Δ*ABC*) mutant produces VPS but no accessory proteins (39); it shows emergent bipolar order near the biofilm-gel interface, but the interior remains disordered. When the surface adhesion proteins are present as in the Δ*rbmA* mutant, the bipolar cell ordering is fully established (**Fig. 4, A** and **B** and Fig. S14), and cells pack more densely at the interface than in the interior (Fig. S2). Interestingly, in the Rg biofilms that produce all matrix components, the bipolar surface order is slightly lower (**Fig. 4B**), and the planar cell ordering at the core is less pronounced (Fig. S15). These results confirm the role of VPS and matrix proteins in mediating cell ordering in embedded biofilms.

**Fig.4.**
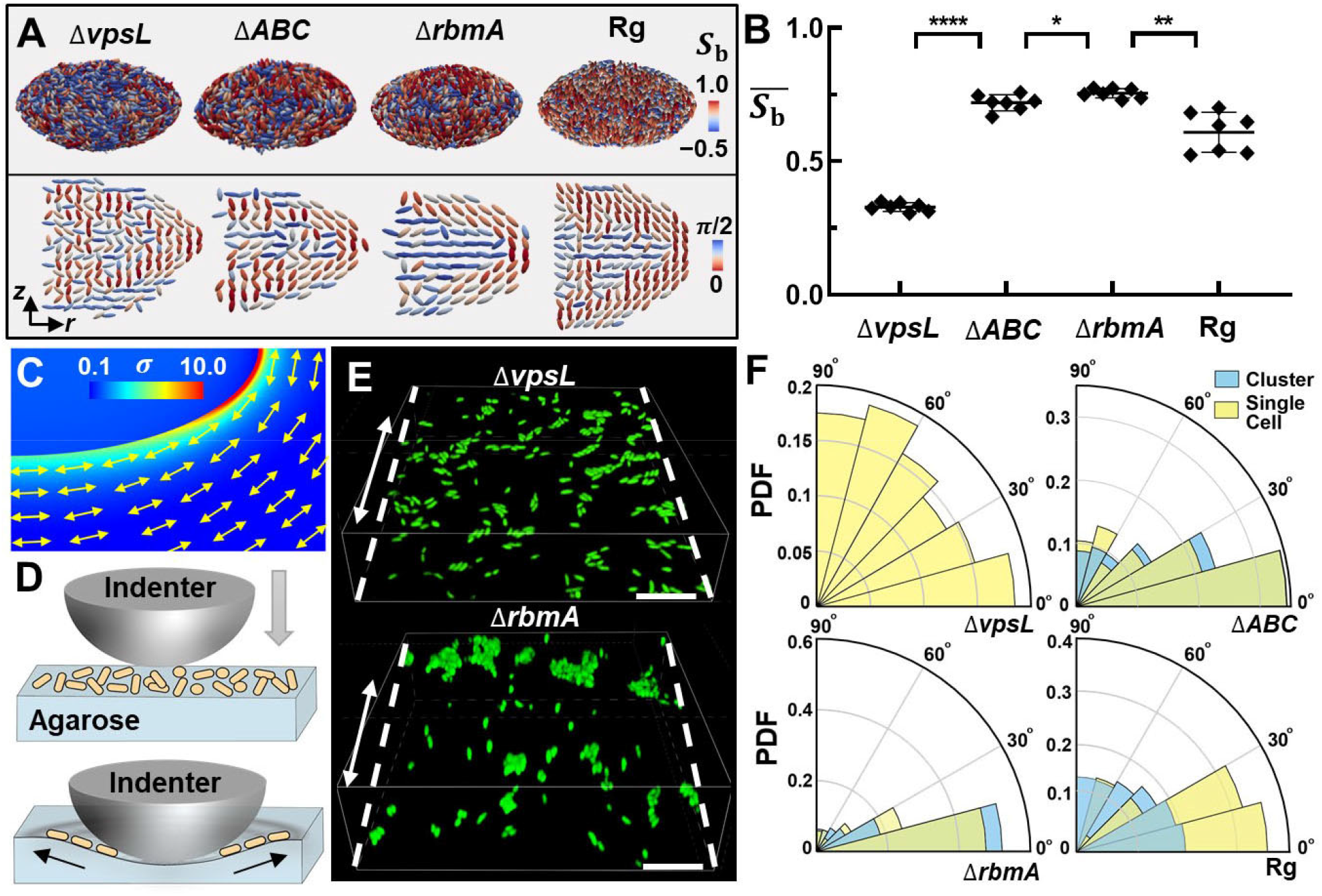
Stress transmission across the biofilm-gel interface by matrix components causes cell ordering. (*A*) Architecture of mature biofilms from different mutants growing inside 2% agarose gels. The same color codes are used as in **Fig. 3D**. Shown from left to right are the embedded colonies from Δ*vpsL*, Δ*rbmA*Δ*bap1*Δ*rbmC* (Δ*ABC*), Δ*rbmA*, and rugose wild-type (Rg) cells. (*B*) Average bipolar ordering 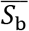 calculated for each mutant biofilm. Unpaired *t*-test is performed for statistical analysis; *P<0.05; **P<0.01; and ****P< 0.0001; error bars indicate SDs (*n* = 7). (*C*) Finite element simulation result for the normalized von Mises stress field (*σ*/*G*_b_) generated by inclusion growth with Δ*V/V*_0_ = 25. Double-headed arrows indicate the local directions of the first principal tensile stress in the gel. (*D*) Schematic illustration of the *in vitro* stress transmission experiment. A spherical indenter generates a radially symmetric deformation field in the agarose gel, whose surface is colonized by randomly oriented cells (yellow cylinders) and small clusters. After indentation, we record the orientation of cells in areas not in physical contact with the indenter. (*E*) Images of single cells and clusters of Δ*vpsL* and Δ*rbmA* mutant cells on the deformed agarose surface stretched along the direction of dashed line. Scale bar: 10 μm. (*F*) Probability distribution function (PDF) of the angles between the axis of cell orientation and the axis of substrate stretching (black arrow in (*D*)) for different mutants. Yellow corresponds to single cells and blue corresponds to cells in clusters.

How do the molecular-scale interactions mediated by the matrix components translate into community-scale organization? A hint is given by finite element simulation (**Fig. 4C**). Due to the volumetric expansion of the biofilm, the gel material near the interface experiences large *tensile* stresses. Importantly, the direction of tensile stresses coincides with the direction of cell alignment. Inspired by how epithelial cells align along the axis of maximum principal stress (40), we hypothesize that bacterial cells at the biofilm-gel interface align with the tensile stresses in the gel, and that this stress transmission depends on the biofilm matrix. To test this hypothesis, we design an *in vitro* indentation experiment to capture the essence of the mechanical response at the biofilm-gel interface (**Fig. 4D**). We use a spherical indenter to deform the agarose substrate, generating radial tensile stress. In this experiment, the bulk compressive stress from the confining environment is absent so that we can focus on how bacteria respond to the tensile stress in the substrate, either as individual cells or as a collective. To this end, immediately after indentation, we measure the orientations of isolated cells and cells in small clusters that are randomly attached to the surface (**Fig. 4E**). We calculate the distribution of the angle between the axis of cell orientation and that of the local tensile stress in the substrate (**Fig. 4F**). We find that Δ*vpsL* cells maintain a random orientation, suggesting that in the absence of VPS, *V. cholerae* cells are not able to respond to external stress. This observation explains the disordered organization of the Δ*vpsL* colony. On the other hand, the Δ*ABC* cells readily align with the tensile stress direction, consistent with the emergence of bipolar order in Δ*ABC* biofilms. Interestingly, this result indicates that VPS alone can transmit stress across the biofilm-gel interface (Fig. S16). Similar reorientation dynamics is observed in the Δ*rbmA* mutant, and the degree of alignment is stronger than that in the Δ*ABC* mutant. This is consistent with the stronger ordering observed in the Δ*rbmA* biofilm.

A distinct pattern is observed for the Rg strain in the indentation experiment. Isolated cells on the gel surface readily align with the tensile stress similar to the Δ*rbmA* or Δ*ABC* mutant. However, this mechanical response diminishes significantly in clusters – Rg cells in clusters are nearly randomly oriented and unresponsive to the tensile stress in the substrate. We infer that the cell-to-cell adhesion conferred by RbmA inhibits the reorientation process (11, 13, 39) – it may be difficult for mutually adhered cells to reorient collectively and transition from the initial random configuration to the bipolarly ordered configuration. The impaired alignment with external stress underlies the decreased bipolar order in the embedded Rg biofilms (Fig. S15). To sum up, the *in vitro* indentation experiment provides direct evidence that *V. cholerae* cells can align with tensile stress in an adjacent elastic substrate, and that this response depends on the matrix components. The bipolar organization of cells in embedded biofilms is therefore a consequence of this alignment to the tangential tensile stress in the gel.

## Discussion

In this study, we use a highly tunable biomechanical system consisting of *V. cholerae* biofilms that develop inside a biocompatible hydrogel to obtain insight into how growing bacterial communities interact with their mechanical environment. While mechanical stress has been implicated to play an important role in surface-attached biofilms (9–15), embedded biofilms face unique mechanical conditions in which they must deform and even damage their environment as they grow. Our study employs new experimental and theoretical approaches to understand this constrained growth process. The key mechano-morphogenesis steps and their dependence on the mechanical environment are studied by combining single-cell live imaging, mutagenesis*, in vitro* rheological testing, theoretical mechanics, and numerical modeling. We reveal that the morphodynamics and internal structure of embedded biofilms crucially depends on the relative stiffnesses of the biofilm and its environment. Biofilms growing in a stiff environment develop oblate shapes and bipolar cellular ordering, which result from the transmission of growth-induced stresses across the biofilm-environment interface. In contrast, biofilms in soft environments feature a spherical shape and random cellular organization.

The morphogenetic principles of confined *V. cholerae* biofilms revealed here are generally applicable to other biofilms and potentially to higher-order organisms developing in mechanically constrained environments. As a proof of principle, we show that embedded colonies formed by the sphere-shaped bacterium *Staphylococcus aureus* (41) (Fig. S17) also exhibit the characteristic oblate shape with an internal organization that is distinct from the morphologies and ordering that emerge in unconfined growth (42). This result with spherical bacteria confirms that the anisotropic biofilm shapes arise not from anisotropic cell shapes but from biofilm-gel interactions.

Our results also suggest new directions for future studies on embedded biofilms, such as their antibiotic resistance. Indeed, bacterial colonies embedded in hydrogels have been shown to display enhanced antibiotic resistance compared to planktonic cultures (41, 43). Our singlecell imaging technique provides quantitative tools to investigate the antibiotic response of individual bacterial cells in this context as demonstrated in Fig. S18, where different degrees of confinement give rise to varying antibiotic responses. Another natural application of the imaging tools is to study bacteria-phage interactions. The sample geometry in this study is widely used in bacteriophage assay in which bacterial colonies are embedded in agar gels, allowed to grow, and subsequently infected with phages (44). We hypothesize that the structural characteristics of a biofilm such as density, cell ordering, and overall morphology, could potentially affect phage infection kinetics, for example by influencing the diffusion dynamics of phages inside an embedded biofilm. Beyond bacterial biofilms, the continuum mechanical theory that we developed for growing inclusions is potentially generalizable to other developmental systems such as growing tumors (21) and organoids (45).

Our findings on confined biofilms invite interesting comparisons with the well-studied case of surface-attached biofilms (9, 11–13). In surface-attached biofilms, cell-to-surface adhesion and cell proliferation jointly lead to a 2D-to-3D transition, initiated by individual cells undergoing a verticalization instability and resulting in mature biofilms with a vertically aligned core and a radially aligned periphery (9, 10, 46, 47). In contrast, when growing under 3D confinement, we find that *V. cholerae* biofilms develop a distinct architecture by changing their overall shape and cell ordering in response to the confinement-induced stress. Towards the late stage of the embedded biofilm growth, cells near the Boojum points do become verticalized with respect to the local interface (**Fig. 3** and Fig. S13). However, this verticalization event has little effect on the global morphology of the embedded biofilms.

Our results suggest that the intricate 3D cell ordering in confined biofilms impinges critically on the stress transmitted through the biofilm-gel interface by the extracellular matrix. An interesting finding is that VPS, while not sufficient to help *V. cholerae* biofilms resist flow in the absence of RbmC/Bap1 (9, 38, 39), is capable of transmitting stress between the gel and the biofilm (**Fig. 4**). Indeed, we have detected a weak but nonnegligible adhesion conferred by VPS in quantitative adhesion measurements using a rheometer (Fig. S16). This residual adhesion, potentially enhanced by the build-up of normal pressure at the biofilm-gel interface, is presumably sufficient for stress transmission. Still, stress transmission is significantly enhanced with the expression of RbmC/Bap1, consistent with their primary function as surface adhesins (37–39). While our results reveal the mechanism underlying cell ordering at the biofilm-gel interface, how the cell ordering at the interface propagates into the biofilm interior is currently unclear. One potentially related observation is the channels percolating the interior of an embedded biofilm (Movie S1, Fig. S19), which are filled with VPS and generally align with the meridians.

An important question remains in the current system as to whether the observed cell ordering can actively drive changes in biofilm morphology, thus affecting the developmental pathways of embedded biofilms. Treating the growing biofilm as a 3D active nematic medium might shed light on this intriguing possibility (11, 48). Specifically, in addition to the two +1 defects on the biofilm surface, we also observe hints of a +1/2 defect ring in the biofilm interior, lying on the principal plane of oblate-shaped biofilms (**Fig. 3C** and **Fig. 4A**). While +1/2 defects have been shown to organize flows in 2D active nematics (49, 50) and to promote the expansion of bacterial colonies (51, 52), their role in 3D growing nematics remains to be explored.

## Methods

### Strains and media

All *V. cholerae* strains used in this study were derivatives of the wild-type *V. cholerae* O1 biovar El Tor strain C6706 and listed in Table S4. The rugose (Rg) strain harbors a missense mutation in the *vpvC* gene (*vpvC*^W240R^) that elevates intracellular c-di-GMP levels (27). The Rg strain forms robust biofilms and thus allows us to focus on the biomechanical mechanisms governing biofilm formation rather than mechanisms involving gene regulation. Additional mutations were genetically engineered into this *V. cholerae* strain using the MuGENT method (53). All strains were grown overnight in LB at 37°C with shaking. Embedded biofilm growth was performed in M9 medium, supplemented with 2 mM MgSO_4_, 100 μM CaCl_2_, and 0.5% glucose.

### Biofilm growth

*V. cholerae* strains were first grown overnight at 37°C in liquid LB with shaking, back-diluted 30-fold, and grown for an additional 2 hours with shaking in M9 medium at 30°C until early exponential phase (OD_600_ = 0.1–0.2). In the meantime, agarose liquid gels were prepared by microwaving M9 medium containing the designated concentrations (0.3%-2.5% (w/v)) of agarose powder and kept warm at 45°C. For time-course imaging, the regrown cultures were diluted to OD_600_ = 0.0002 with the warm agarose liquid. Subsequently, 50 μL of the inoculated gels was added to wells of 96-well plates with #1.5 coverslip bottom (MatTek). The gels were allowed to cool down and solidify for 10 minutes at room temperature, and subsequently, 100 μL of fresh M9 medium was added on top of the gel. The low initial inoculation density enabled isolated biofilm clusters to grow inside the gel. The locations of the founder cells were identified, and imaging began on the microscope stage two hours after inoculation. A stage-top incubator (Tokai Hit) was used to keep the sample at 30°C. For endpoint imaging, the inoculated samples were incubated at 30°C for the designated duration before imaging. Biofilms grown in this geometry can grow up to 3 days, after which significant cell death occurs even with the replenishment of fresh media (Fig. S20).

### Microscopy

Confocal images were acquired with a Yokogawa CSU-W1 confocal spinning disk unit mounted on a Nikon Ti2-E inverted microscope with a Perfect Focus System, using a 60× water (N.A. = 1.2, for single-cell imaging) or a 20× multi-immersion objective (N.A. = 0.75, for morphological analysis) for different experiments. Cells expressing mNeonGreen were excited with a 488 nm laser, and the fluorescent signal was captured by a sCMOS camera (Photometrics Prime BSI). For time-lapse imaging, to avoid evaporation, immersion oil with a refractive index of 1.3350 (Cargille) was used instead of water and samples were kept in a stage-top incubator (Tokai Hit). To obtain sufficient magnification in single-cell imaging for automated cell segmentation, a 1.5× post-magnification lens was placed between the CSU unit and the microscope side port. The magnification was 72.2 nm per pixel in the *x–y* plane, with a 210 nm step size in the *z* direction (Fig. S1). The point spread function (PSF) of the system was measured under the same imaging conditions (with a 50 nm *z-*step size) using 100 nm fluorescent TetraSpeck microspheres (Invitrogen) embedded in 1% agarose gels. The time difference between each image acquisition was 30 minutes to one hour, and the total acquisition time was 18 hours for each experiment. To further decrease phototoxicity to the cells, an adaptive *z* range was used. For morphological characterization, a 20× water objective plus 1.5× post magnification was used. The magnification was 660 nm per pixel in the *x–y* plane, with a 1320 nm step size in the *z* direction. All image acquisitions were automated using Nikon Element software. All images presented in this work are raw images from this step, rendered by Nikon Element software.

### Statistical methods

Error bars correspond to standard deviations from measurements taken from distinct samples. Standard *t*-tests were used to compare treatment groups and are indicated in each figure legend. Tests were always two-tailed and unpaired as demanded by the details of the experimental design. All statistical analyses were performed using GraphPad Prism software.

## Acknowledgments

We thank Drs. A. Mashruwala and N. Høyland-Kroghsbo for their help in the initial experiments. We thank Drs S. Mao, A. Crosby, E. Dufrense, and J. S. Tai for helpful discussions. T.C. acknowledges the support of Dr. Timothy B. Bentley, Office of Naval Research Program Manager, under award number N00014-20-1-2561, and support from the National Science Foundation under award number 1942016. R.A. acknowledges support from the Human Frontier Science Program LT000475/2018-C. J.Y. holds a Career Award at the Scientific Interface from the Burroughs Welcome Fund.

## Author contributions

Q.Z. and J.Y. designed and performed the experiments. J.L., M.K., and T.C. performed modeling. J.Y. performed strain cloning. Q.Z., H.L., J.N., and J.Y. analyzed the data. Q.Z., J.L., R.A., T.C., M.K., and J.Y. wrote the paper.

## Supplementary Information

is available.

## Data and materials availability

Matlab codes for single-cell segmentation are available online at Github: https://github.com/Haoran-Lu/Segmentation_3D-processing/releases/tag/v1.0. Other data are available upon request.

