## Supporting Information for "Morphogenesis and cell ordering in confined bacterial biofilms"

**This PDF file includes:**

Materials and Methods
Figs. S1 to S20
Tables S1 to S4
Captions for Movies S1 to S4
References

**Other Supplementary Materials for this manuscript include the following:**
Movies S1 to S4(.mov)

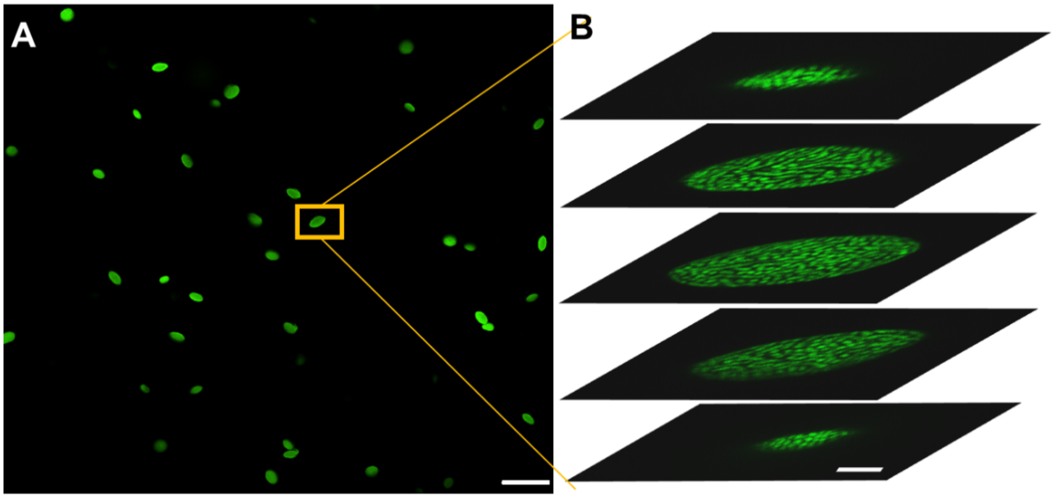


**Fig. S1. Embedded biofilm clusters imaged with spinning disc confocal microscopy.** (*A*) Representative large-scale view of *V. cholerae* biofilms embedded in 2% agarose gel imaged with 20× water objective, showing random orientations of the biofilm clusters. Scale bar: 100 µm. (*B*) Zoom-in images at five different *z* planes of a biofilm cluster for single-cell analysis, imaged with a 60× water objective with 1.5× post-magnification. Scale bar: 5 µm.

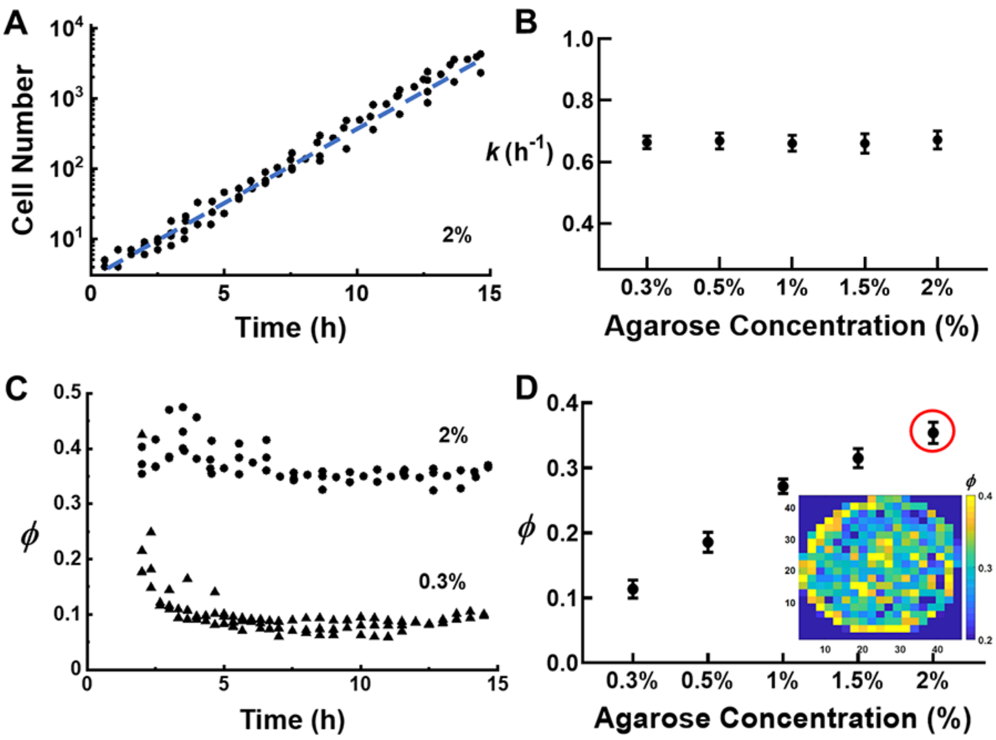


**Fig. S2. Structural properties of biofilms grown inside gels solidified with different agarose concentrations.** (*A*) Growth curves of individual Δ*rbmA* biofilm clusters in 2% agarose gel, showing the exponential growth of embedded biofilm at a constant growth rate $k$ ($n$ = 3). (*B*) The growth rate $k$ of Δ*rbmA* biofilms versus agarose concentration (error bars correspond to SDs). No statistical difference is found between gels of different agarose concentrations, suggesting that the mechanical constraints relevant in this study do not affect cell growth. (*C*) Time evolution of cell density quantified by the volume fraction $\phi$, i.e. the fraction of biofilm space occupied by cells (1) for the Δ*rbmA* biofilm clusters growing in 0.3% and 2% agarose gels ($n$ = 3). (*D*) $\phi$ versus gel concentration (error bars correspond to SDs, $n$ = 5). Inset shows the spatial distributions of $\phi$ in a representative biofilm embedded in 2% agarose. The density of the outer layer is higher than that of the inner layer, which is presumably caused by denser cellular packing at the biofilm-gel interface.

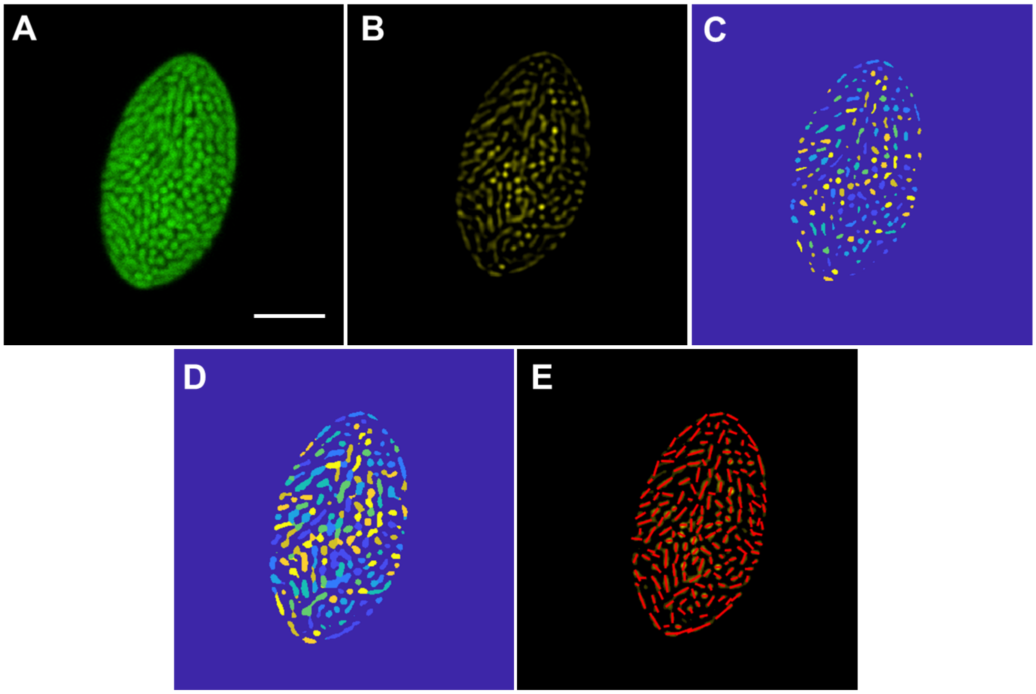


**Fig. S3. Image analysis procedure.** (*A*) Raw cross-sectional image of the middle layer of an embedded biofilm formed by Δ*rbmA* mutant cells. Scale bar: 5 µm. (*B-D*) The corresponding image after deconvolution (*B*), after segmentation with the adaptive thresholding algorithm (*C*), and after a volume-restoration algorithm (*D*). Cell identity as indicated by different colors remains the same in C and D while the volume of each cell was restored to the correct value. (*E*) After segmentation and volume restoration, the orientation of each cell was inferred. A red line with a set length was drawn for each segmented cell, which was subsequently projected onto a 2D plane. A long red line in the projected plane represents a cell lying approximately parallel to the imaging plane while short lines represent vertical cells.

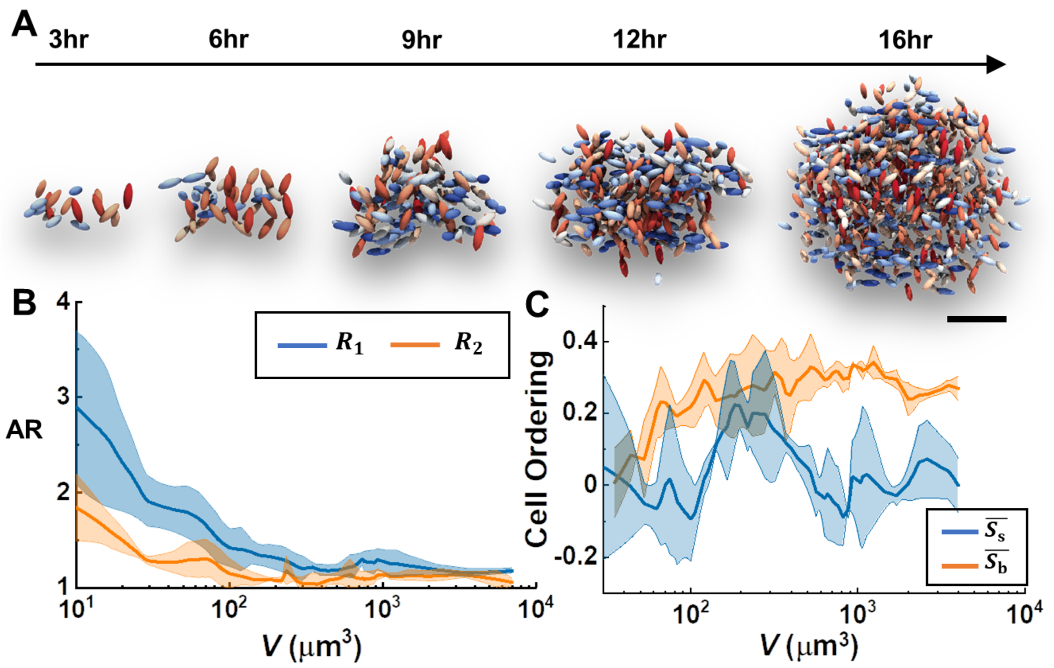


**Fig. S4. Biofilm growth inside soft gels.** (*A*) Single-cell 3D reconstructed images for a biofilm cluster growing inside 0.3% agarose gel. Scale bar: 5 μm. (*B*) Evolution of biofilm morphology characterized by aspect ratios (ARs) including $R_1$ (blue) and $R_2$ (orange) inside 0.3% agarose gel as a function of biofilm volume $V$. Center lines correspond to means; widths of the shaded areas correspond to SDs ($n$ = 5). (*C*) Evolution of cell ordering with biofilm volume $V$, quantified by the shell ordering $\overline{S_s}$ and bipolar ordering $\overline{S_b}$ in the outmost layers. Center lines correspond to means; widths of the shaded areas correspond to SDs ($n$ = 5).

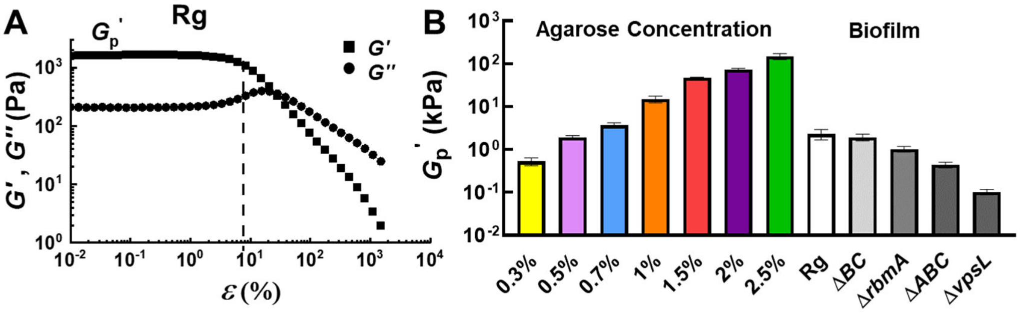


**Fig. S5. Rheological characterization of *V. cholerae* biofilms and agarose gels.** (*A*) Representative storage modulus $G'$ and loss modulus $G''$ curves as a function of the amplitude of oscillatory shear strain $\varepsilon$ measured for the Rg biofilm. From the $G'(\varepsilon)$ curves, the storage modulus plateau value $G_p'$ was extracted for each biofilm. (*B*) Measured $G_p'$ value of agarose gels with different concentrations and of different mutant biofilms. All biofilms were grown for two days on M9 medium solidified with 1% agar. Error bars correspond to SDs ($n = 3$).

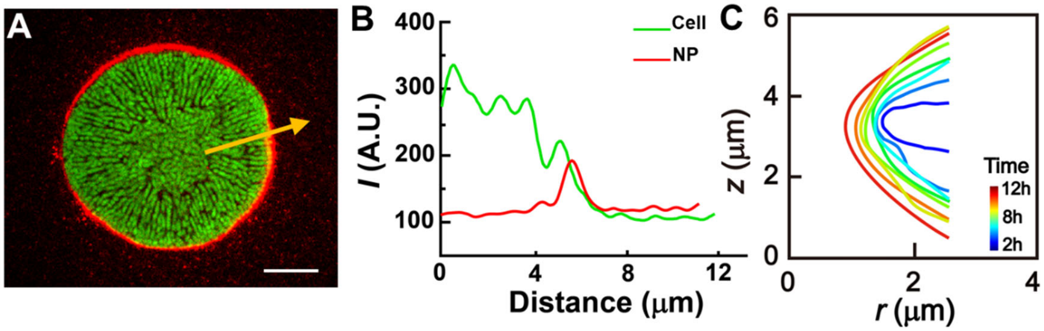


**Fig. S6. Evidence for the absence of crack in gel.** (*A*) Image of a biofilm growing in 2% agarose gel with embedded nanoparticles (diameter = 40 nm, red). After biofilm growth, nanoparticles were excluded from the biofilm region and therefore accumulate at the edge of the embedded biofilm; no nanoparticle-excluding zone was observed, indicating the absence of macroscopic cracks. Scale bar: 5 μm. (*B*) Fluorescence intensity $I$ versus radial distance $r$ of biofilm image in A along the radial direction for the cell channel (green) and the nanoparticle (NP) channel (red), respectively. (*C*) Evolution of biofilm contour in the $x$-$z$ plane, showing that the curvature at the biofilm edge increases with biofilm growth. Curves are shifted for clarity and color-coded according to time. In the hypothetical crack scenario, the shape of the biofilm edge should instead stay constant as the crack propagates (2).

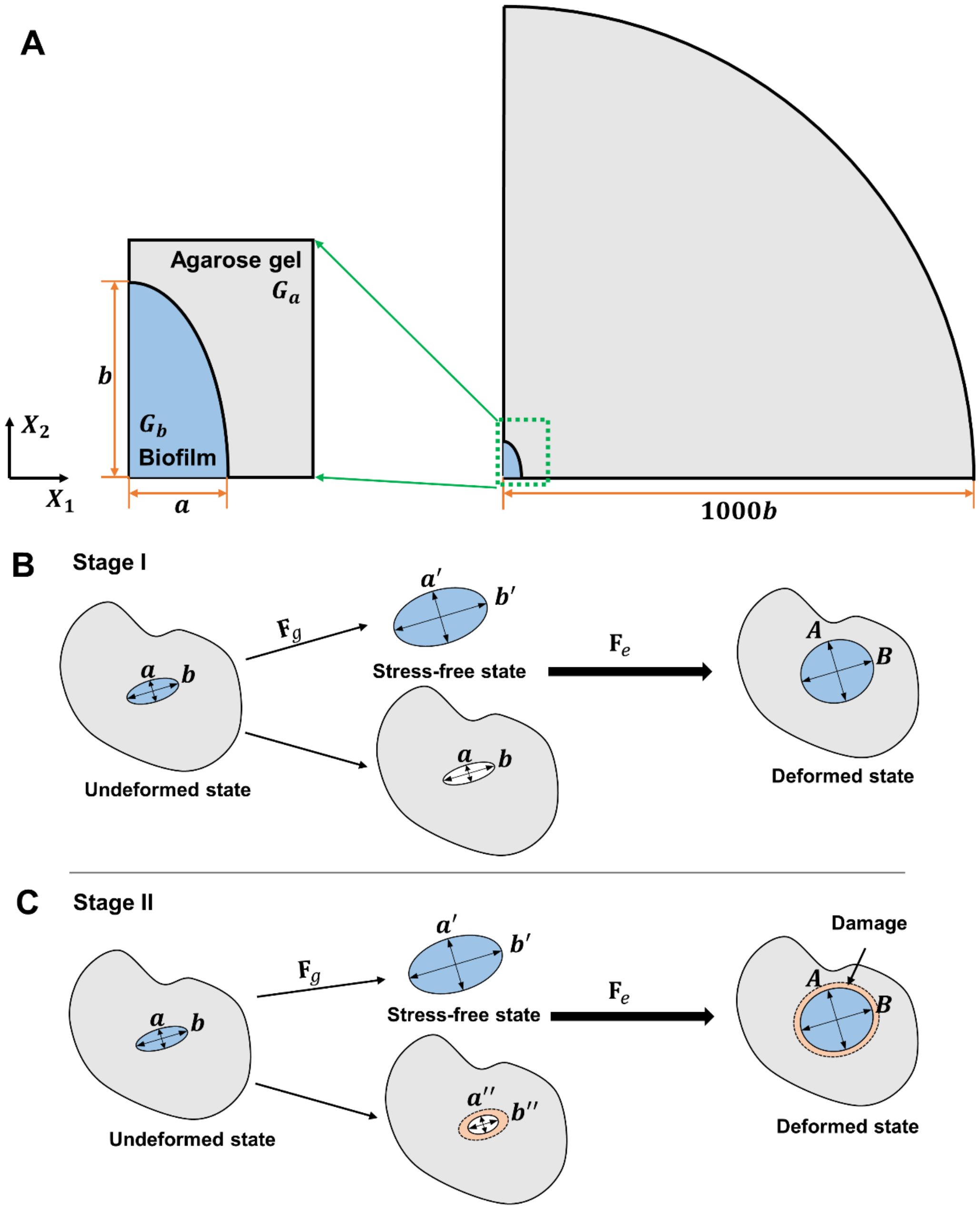


**Fig. S7. Summary of the continuum modeling approach.** (*A*) Computational model configuration. (*B*) Stage I (purely elastic deformation), the agarose gel undergoes small to moderate strains and the deformations are purely elastic; correspondingly, in the simulation model, the void retains its stress-free shape. At each volume expansion ratio $\Delta V/V_0$ the biofilm chooses its stress-free shape in a way that minimizes the total elastic energy in the system. (*C*) Stage II (damage transition). The damage in the gel is modeled by varying the effective shape of the void, while the stress-free shape of the biofilm and the void are assumed to be the same, thus avoiding any shape mismatches. The optimal shape of biofilm at each volume expansion ratio $\Delta V/V_0$ is obtained by minimizing the total stored energy in the system via changing the stress-free shape of the biofilm.

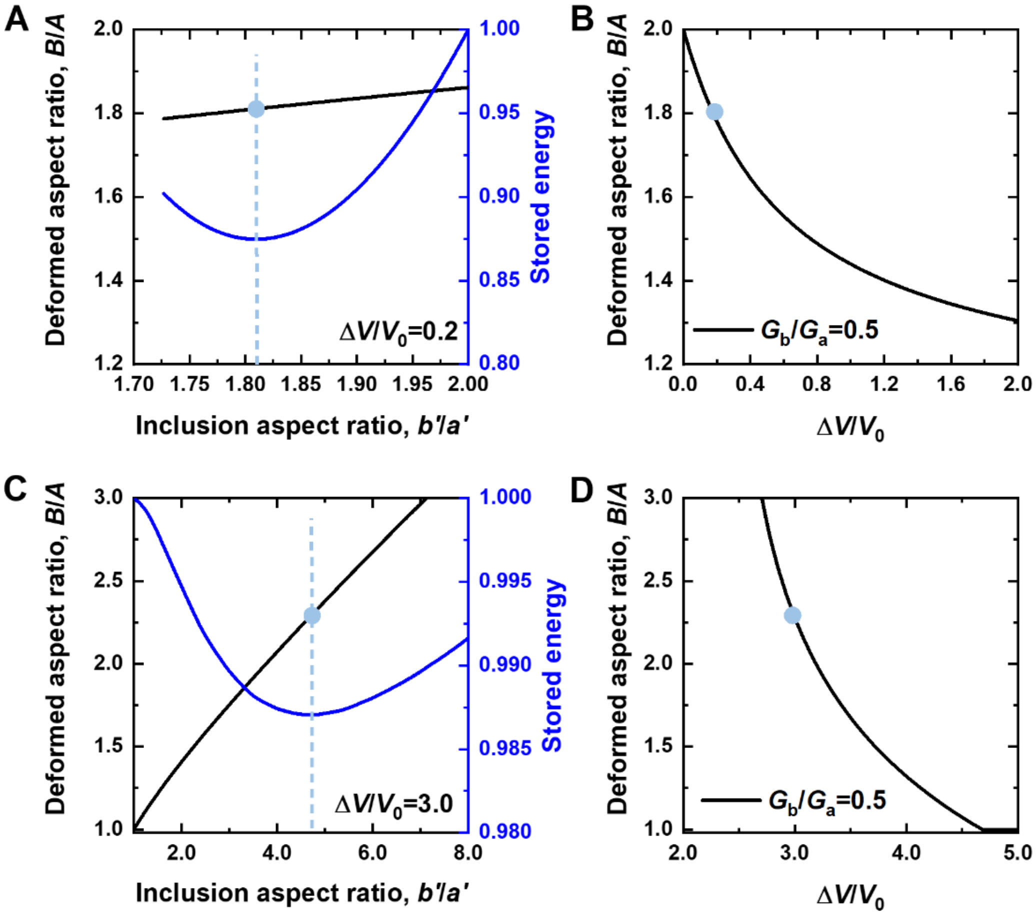


**Fig. S8. Typical results from mechanical energy minimization in modeling.** (*A-B*) Purely elastic deformation (Stage I): An example is shown to demonstrate how the optimal inclusion shape at a particular volume expansion ratio is obtained. Plotted are the aspect ratio of the inclusion in the deformed state (*B*/*A*, left $y$-axis) and the corresponding stored mechanical energy (right $y$-axis) versus inclusion aspect ratio in the stress-free state, $b'/a'$. The dashed line corresponds to the position at which the stored energy reaches minimum, from which the optimal *B*/*A* was obtained (blue dot). The stored energy is normalized by the case with isotropic volume expansion (i.e. $\alpha = \beta$, meaning that $b'/a' = b/a = 2.0$). (*B*) The optimal *B*/*A* as a function of volume expansion $\Delta V/V_0$ for the case of $G_\mathrm{b}/G_\mathrm{a} = 0.5$. (*C-D*) The same calculation was repeated in Stage II (the damage transition). The stored energy is normalized by the case with circular inclusion (i.e. $b'/a' = 1$).

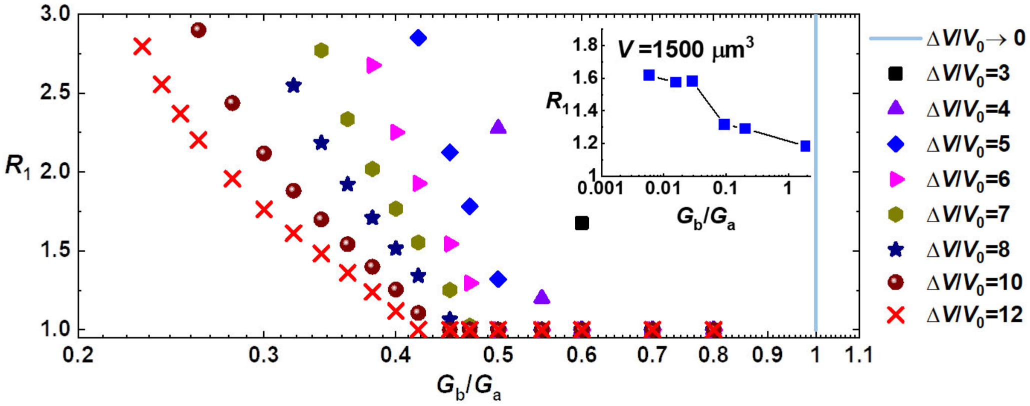


**Fig. S9. Qualitative comparison of the modeled and the experimentally measured biofilm aspect ratio $R_1$.** Shown in the main panel are the predicted $R_1$ values calculated for different values of the volume expansion ratio $\Delta V/V_0$ at the onset of damage (Stage II) as a function of the stiffness contrasts $G_b/G_a$ in the theoretical model. Inset shows the experimentally measured $R_1$ as a function of $G_b/G_a$ at $V$ = 1500 μm$^3$. In the experiment, $G_b/G_a$ values are tuned by modifying the biofilm stiffness, while fixing the agarose concentration to 2%. For a given $\Delta V/V_0$, both experiment and simulation show that $R_1$ decreases with increasing stiffness contrast $G_b/G_a$.

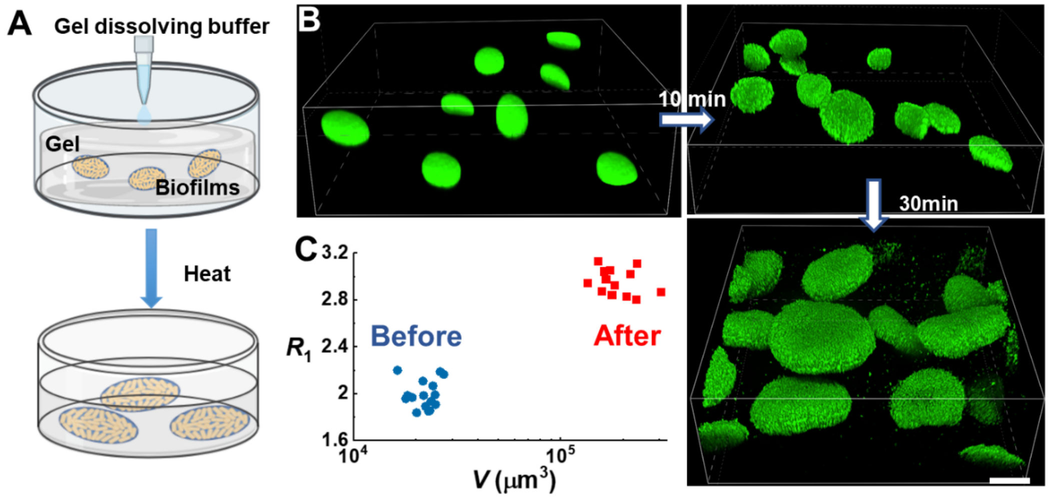


**Fig. S10. Mature biofilms swell and change shape after the confinement is released.** (*A*) Schematic illustration of the agarose dissolution experiment to release mature embedded biofilms from confinement. (*B*) Representative biofilm images taken at different time points after adding gel dissolving buffer and heating at 40°C. Scale bar, 40 μm. (*C*) Morphological change of embedded biofilms before and after dissolving agarose, quantified by their $R_1$ and $V$. Swollen biofilms have a larger volume and a higher aspect ratio. This observation is consistent with our model: the inclusion in the hypothetical stress-free state is more anisotropic than the embedded state (Fig. S7).

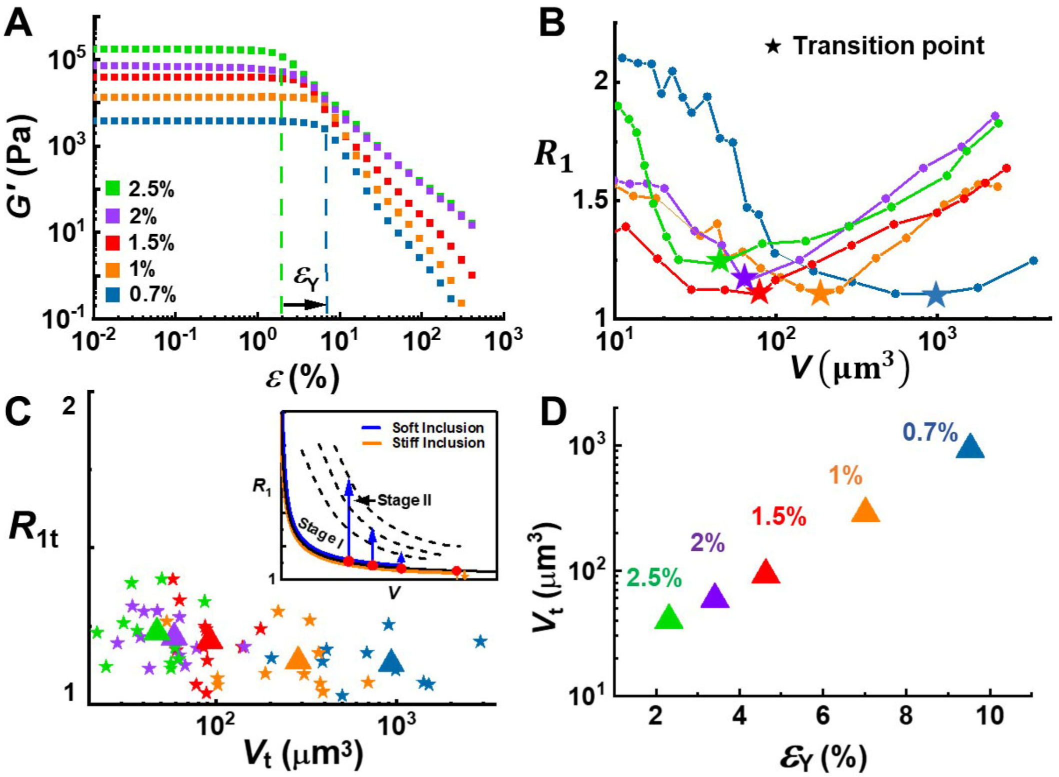


**Fig. S11. Biofilm shape evolution depends on gel failure mechanics.** (*A*) Representative storage modulus $G'(\varepsilon)$ curves measured for gels solidified with various agarose concentrations. Yield strain $\varepsilon_Y$ (indicated by the vertical dotted line) was extracted for each gel concentration. (*B*) Evolution of $R_1$ as a function of volume $V$ for biofilms growing inside agarose gels with various concentrations indicated by their color. Stars correspond to the minimum on the $V$-$R_1$ plot, which define the transition biovolume $V_t$ and the transition aspect ratio $R_{1t}$. (*C*) $R_{1t}$ versus $V_t$ measured for biofilms growing in gels solidified with different agarose concentrations. Each point was measured from a single time-lapsed movie of a growing biofilm cluster. Triangles indicate the average. Inset shows a schematic explanation of how the Stage I to Stage II transition happens at different stiffness contrast. Specifically, under all conditions, biofilms follow a universal pathway in Stage I with a monotonically decreasing $R_1$ versus $V$. The yield strain of the gel determines when the transition takes place and the longer the system stays in Stage I, the lower the transition aspect ratio $R_{1t}$. (*D*) Average volume at transition point $V_t$ for biofilms grown in agarose gels with the designated concentrations, as a function of the corresponding yield strain $\varepsilon_Y$ of the gel. $V_t$ scales monotonically with $\varepsilon_Y$, which is consistent with the theoretical model (**Fig. 2E**).

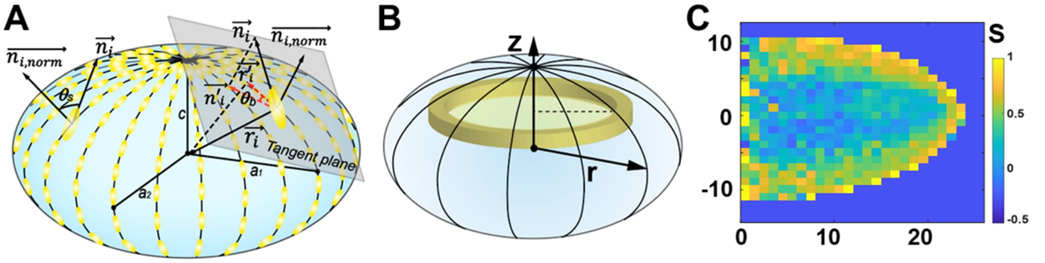


**Fig. S12. Definition and analysis of order parameters.** (*A*) Schematic illustration of the definition for the shell order parameter $S_s$ and bipolar order parameter $S_b$. See Methods for details. (*B*) Schematic illustration of the 2D azimuthally averaged direction and order parameter of cells, projected to the (*r*, *z*) plane. (*C*) Representative measurement of the cell ordering inside biofilms grown in 2% gels. The cell ordering was quantified through a ***Q***-tensor based method (see Methods). Color denotes the scalar order parameter *S*. Only the cells in the periphery show significant nematic ordering.

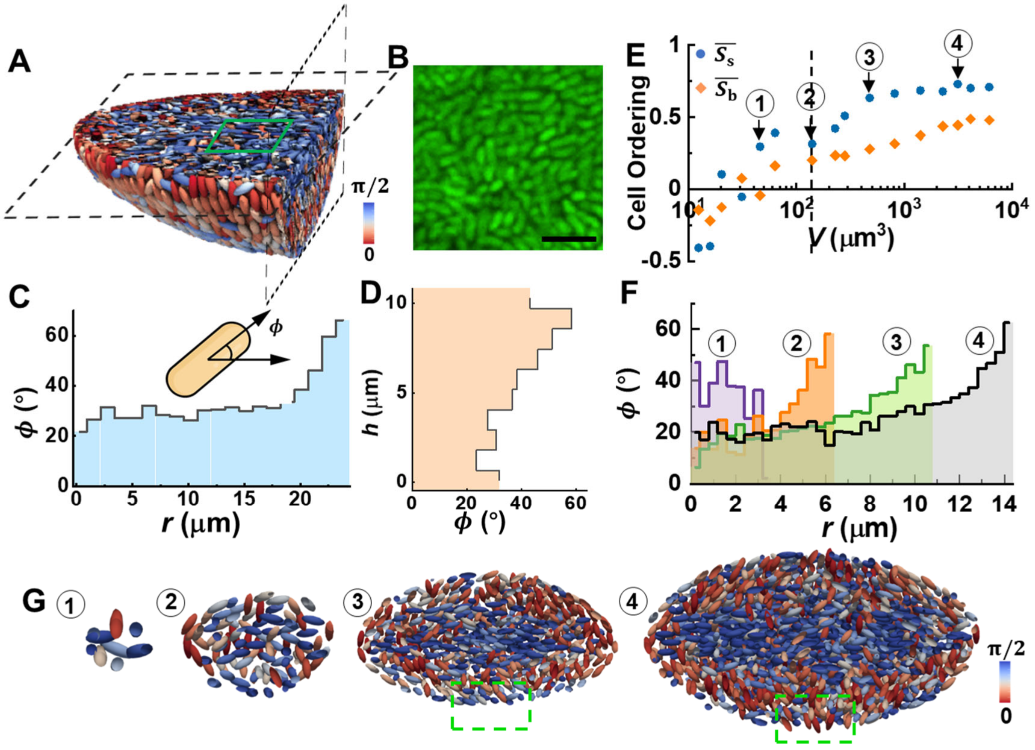


**Fig. S13. Internal architectural transition of embedded biofilms.** (*A*) Reconstructed internal architecture of an embedded biofilm in 2% agarose. Both the horizontal and vertical cross-sections are shown. Cells are color-coded according to their orientations. (*B*) Zoomed-in image of cells at the biofilm core highlighted in A. Scale bar: 5 μm. (*C*) Averaged angle $\phi$ between a cell and the principal plane plotted against *r*, radial distance from the center. (*D*) Averaged angle $\phi$ of cells in the central vertical cylinder with $r'$=0.1*r*, plotted against the vertical coordinate *h*. (*E*) Evolution of cell ordering with biofilm volume *V*, quantified by the shell ordering $\overline{S_s}$ and bipolar ordering $\overline{S_b}$ in the outmost cell layers. (*F*) Representative internal structural evolution, characterized by the average angle $\phi$ as a function of the radial coordinate *r* at various time points. Cells at the biofilm core tend to align at an average angle of ~20° with the principal plane as the biofilm enters Stage II. (*G*) Evolution of cross-sectional view of a reconstructed biofilm growing in 2% agarose. Cells at the pole (green highlighted area) verticalize with respect to the local interface at the late stage of biofilm development. This observation is consistent with the peak in panel D.

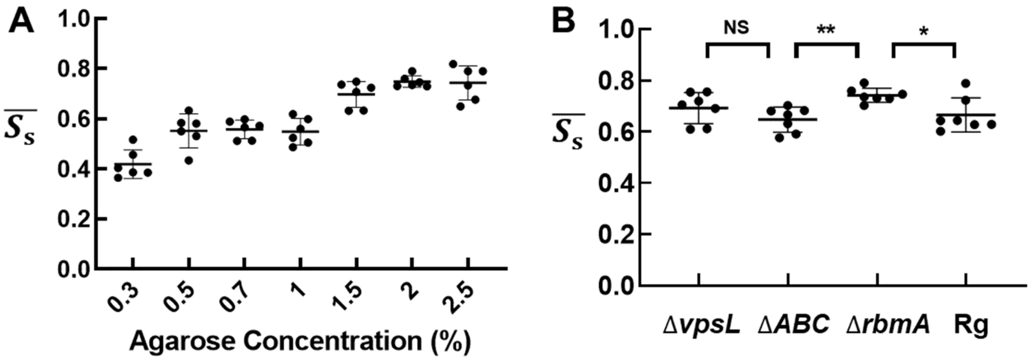


**Fig. S14. Average shell order parameter $\overline{S_s}$ of mature biofilms.** (*A*) $\overline{S_s}$ for biofilms grown in gels solidified with agarose with the designated concentrations. Error bars correspond to SDs (*n* = 6). (*B*) $\overline{S_s}$ calculated for each mutant biofilm in 2% agarose gel. Unpaired *t*-test is performed for statistical analysis; NS denotes not significant; *P< 0.05; and **P< 0.01; Error bars indicate SDs (*n* = 7).

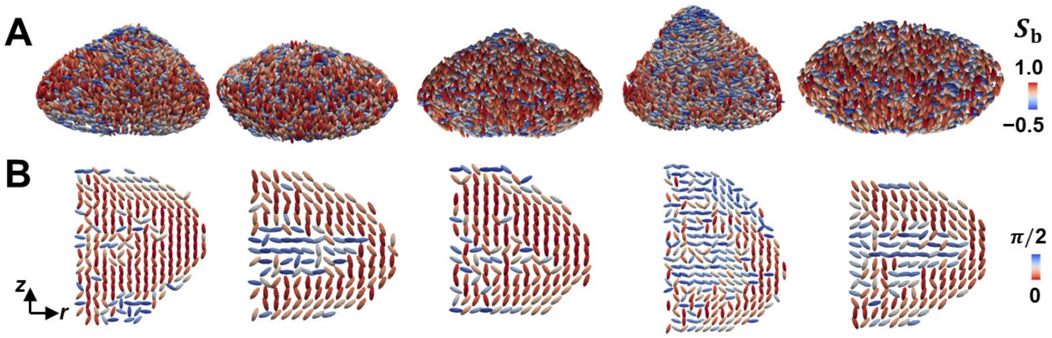


**Fig. S15. Various morphologies and internal architectures of mature Rg biofilms.** (*A*) 3D reconstructed mature Rg biofilms growing inside 2% gel with various morphologies. Cells are color-coded according to $S_b$. (*B*) 2D azimuthally averaged cell orientation inside Rg biofilms projected to the ($r$, $z$) plane, characterized by the angle between the cell director and the $z$ axis. The bipolar surface order $\overline{S_b}$ and the cell ordering at the core of Rg biofilms vary with the biofilm morphology; both are less pronounced compared to the case of Δ*rbmA* biofilms.

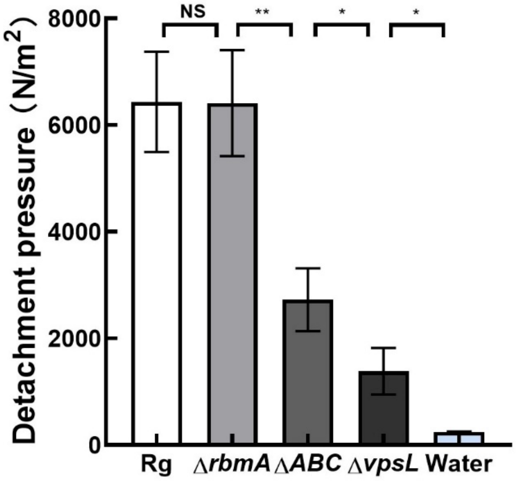


**Fig. S16. Quantification of adhesion strength of *V. cholerae* biofilms with foreign surfaces.** To measure biofilm adhesion strength, we used a modified procedure from the literature (3). Briefly, biofilms from different mutants were first sandwiched between two parallel glass plates separated by 0.3 mm; subsequently, the top plate is gradually lifted, during which the normal force was recorded. We used the peak pressure needed to detach a biofilm from the plate as a quantification of cell-to-substrate adhesion. The detachment pressure of Rg and Δ*rbmA* biofilms are high and comparable to each other, due to the presence of the cell-to-surface adhesion protein RbmC/Bap1. In contrast, the Δ*ABC* mutant biofilm possesses a detachment pressure lower than that of Rg or Δ*rbmA* biofilms but higher than that of Δ*vpsL*. This quantitative measurement reveals small but non-negligible adhesion conferred by VPS even in the absence of RbmC/Bap1, which might be sufficient to transmit stresses across the biofilm-gel interface and cause bipolar cell alignment (Fig. 4B and Fig. 4F). Unpaired *t*-test is performed for statistical analysis; NS denotes not significant; *P< 0.05; and **P< 0.01; Error bars correspond to SDs ($n$ = 3).

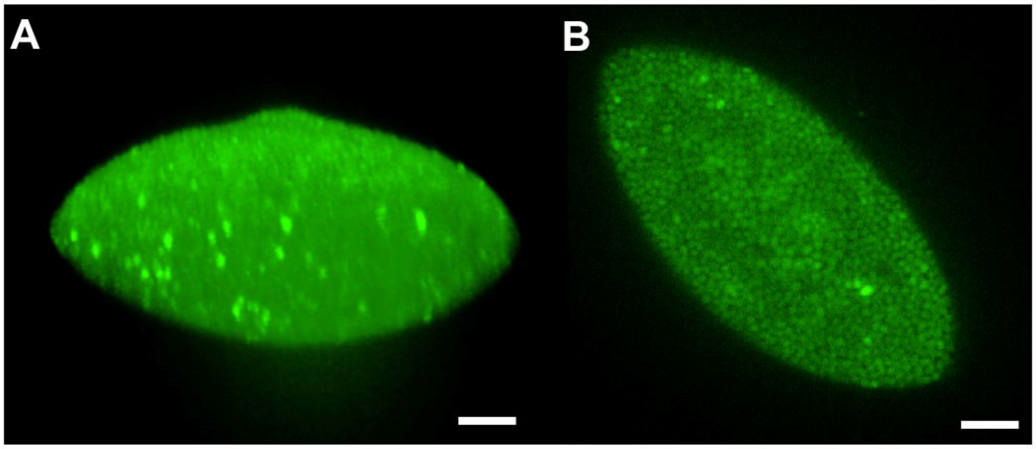


**Fig. S17. Single-cell imaging of embedded colonies from cocci.** (*A*) 3D view and (*B*) cross-sectional image of a *Staphylococcus aureus* (4) cluster embedded in a 2% agarose gel. Similar to rod-shaped bacteria, embedded cocci colonies also acquire an oblate shape under mechanical confinement. This result suggests the generality of our mechanical analysis and indicates that the overall colony shape is independent of the shape of the constituent cells. Compared with biofilm structures grown freely on solid substrates (5), *S. aureus* cells in a confining environment are more densely packed, and they display a certain degree of hexagonal ordering. Scale bar: 5 μm.

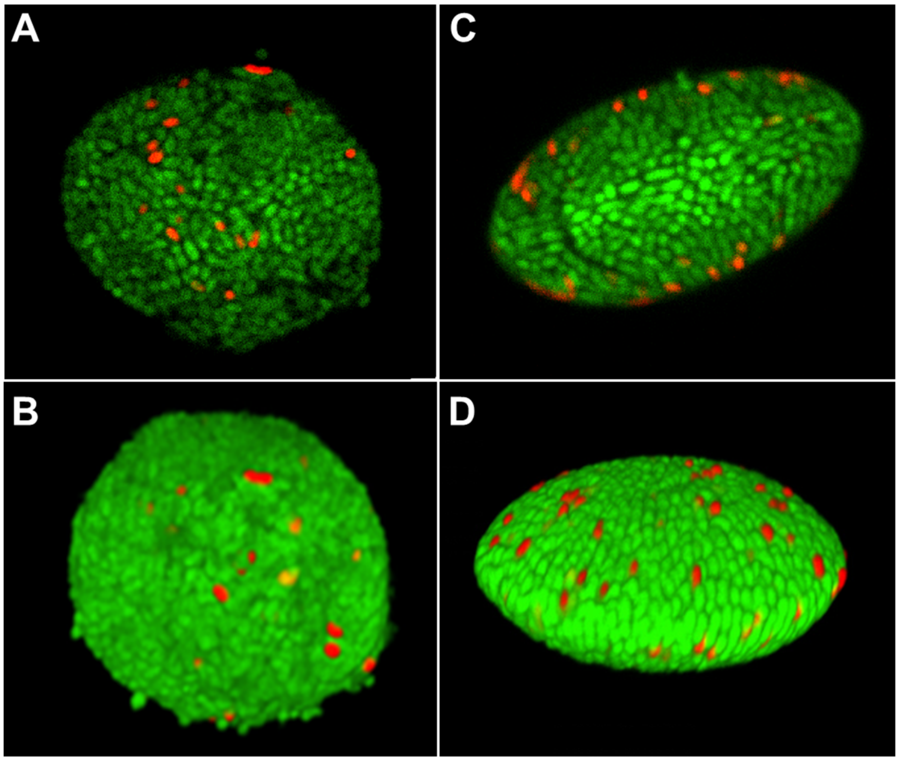


**Fig. S18. Mechanical confinement affects the antibiotic response of cells in embedded biofilms.** (*A-B*) Cross-sectional view (*A*) and 3D view (*B*) of a Δ*rbmA* biofilm with cells constitutively expressing mNeonGreen grown in 0.3% agarose, after antibiotic (tetracycline) treatment. (*C-D*) Cross-sectional view (*C*) and 3D view (*D*) of a biofilm grown in 2% agarose after antibiotic (tetracycline) treatment. Red signal corresponds to dead cells. Cell death occurs primarily at the periphery of embedded biofilms in 2% agarose gel, whereas dead cells are found throughout the biofilms in 0.3% agarose gel.

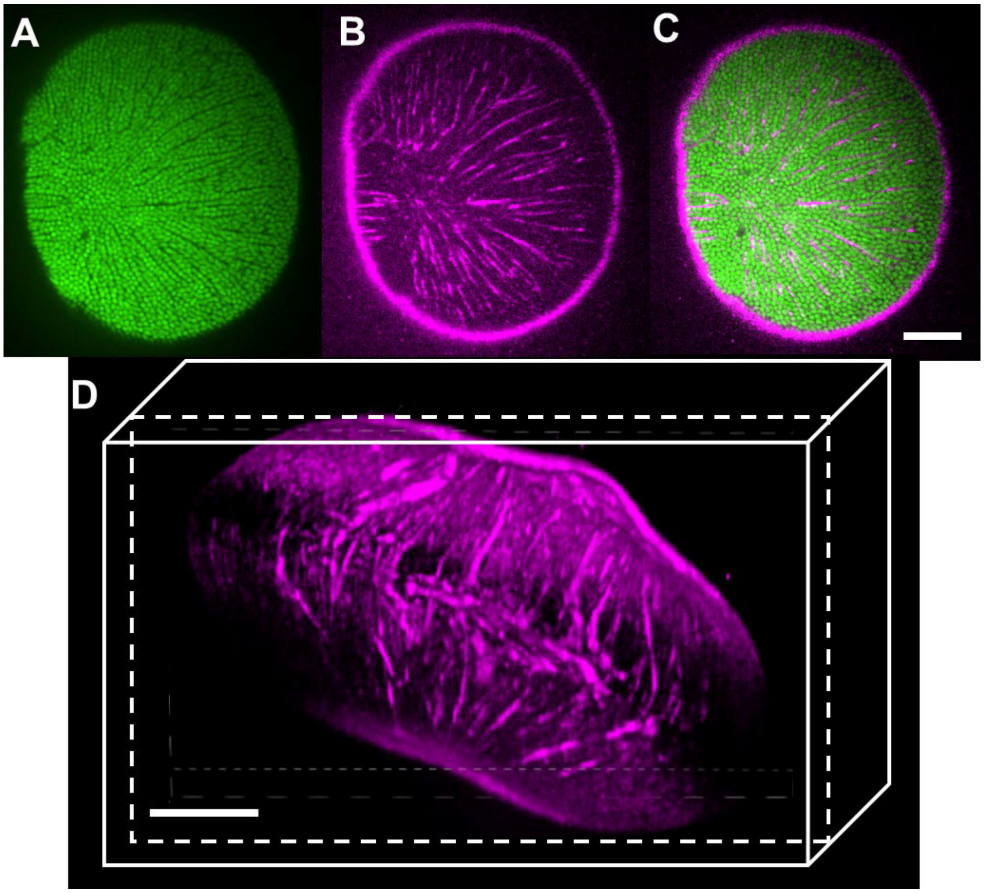


**Fig. S19. Spatial distribution of VPS in an embedded biofilm.** (*A*) A Δ*rbmA* mutant embedded biofilm with cells constitutively expressing mNeonGreen. Channels are visible between densely packed cells. (*B*) VPS is stained with wheat germ agglutinin (WGA) conjugated with Alexa Fluor 647 (6). (*C*) An overlaid image of cells and VPS staining. (D) 3D view of the spatial distribution of VPS signal. Scale bars: 10 μm. This result shows a complete overlap between the location of the channels and the VPS signal. In other words, embedded biofilms possess a network of intra-biofilm channels filled with VPS.

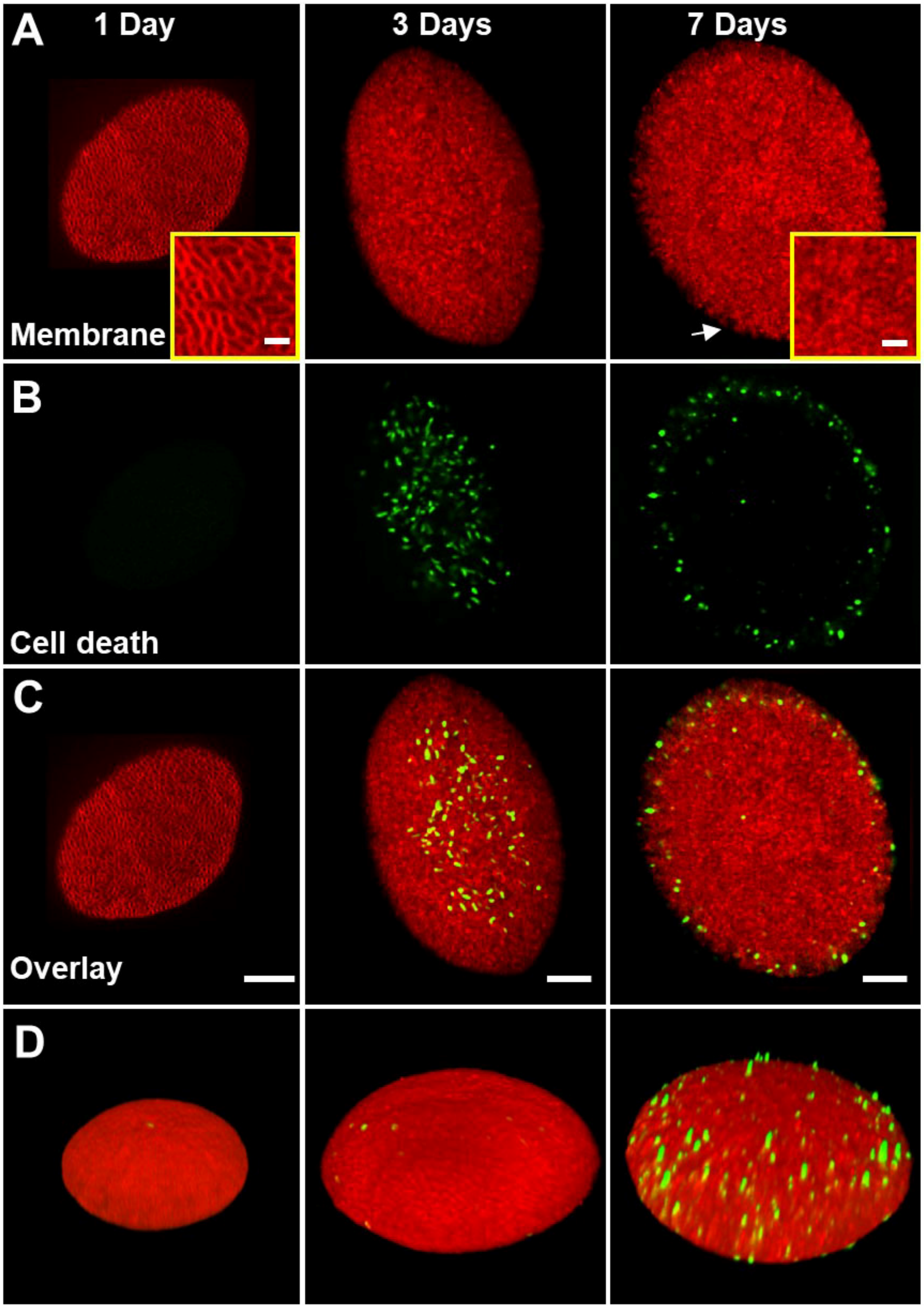


**Fig. S20. Embedded biofilms with longer incubation times.** Representative cross-sectional images of 1-day-old (*Left*), 3-day-old (*Middle*) and 7-day-old (*Right*) embedded biofilms stained with FM 4–64 for cell membranes (*A*) and SytoX Green (7) for cell death (*B*). Insets show the zoom-in images of membrane staining, in which intact membranes with cell contours are observed at day 1 and deteriorated cell membranes are observed at day 7, respectively. (*C*) and (*D*) show the overlaid images and 3D views, respectively. Scale bars: 10 μm. During the long-term experiments, the growth medium was replaced every 2 days with fresh medium. The embedded biofilms stop growing after 3 days, and they retain an oblate shape. For 1-day-old biofilms, cells are all alive and have intact membranes. For 3-day-old biofilms, cell death is localized at the interior of biofilms as shown by the enhanced signal of the SytoX Green, presumably due to lower nutrient availability. For 7-day-old biofilms, most of the cells are dead and cells at the center are disintegrated as shown by the diffuse FM 4-64 signal.

**Table S1. Measured gel material properties and biofilm shape transition volume for gels solidified with different agarose concentrations.**

| Agarose Conc. | $G_a$ (kPa) | Parameters $\varepsilon_Y$ (%) | $\sigma_Y$ (kPa) | $V_t$ ($\mu m^3$) |
|---|---|---|---|---|
| 0.3% | $5.3 \pm 1.1\times10^{-1}$ | $2.3 \pm 0.5\times10^{1}$ | $1.5 \pm 0.3\times10^{-1}$ | NA |
| 0.5% | $1.7 \pm 0.2$ | $1.4 \pm 0.3\times10^{1}$ | $3.4 \pm 0.3\times10^{-1}$ | NA |
| 0.7% | $3.7 \pm 0.6$ | $9.5 \pm 2.0$ | $3.6 \pm 0.5\times10^{-1}$ | $9.3\times10^{2}$ |
| 1% | $1.1 \pm 0.3\times10^{1}$ | $7.0 \pm 1.3$ | $5.3 \pm 0.2\times10^{-1}$ | $2.8\times10^{1}$ |
| 1.5% | $3.6 \pm 0.1\times10^{1}$ | $4.6 \pm 1.0$ | $1.3 \pm 0.1$ | $9.2\times10^{1}$ |
| 2% | $6.5 \pm 0.5\times10^{1}$ | $3.4 \pm 1.1$ | $1.9 \pm 0.2$ | $5.9\times10^{1}$ |
| 2.5% | $1.5 \pm 0.2\times10^{2}$ | $2.3 \pm 0.9$ | $3.1 \pm 0.1$ | $4.2\times10^{1}$ |

**Table S2. Measured material properties of *V. cholerae* biofilms.**

| ***V. cholerae*** **Strain** | **Rg** | **Δ*bap1* Δ*rbmC*** | **Δ*rbmA*** | **Δ*rbmA* Δ*bap1* Δ*rbmC*** | **Δ*vpsL*** |
|---|---|---|---|---|---|
| $G_b$ **(kPa)** | 2.3±0.6 | 1.7±0.4 | 1.0±0.2 | $4.4\pm0.7\times10^{-1}$ | $1.0\pm0.2\times10^{-1}$ |

**Table S3. Fracture scaling analysis for biofilm growth in different gels.**

| Agarose Conc. | $G_a$ (kPa) | Fracture energy $\Gamma$ (J/m$^2$) | Elasto-adhesive length $\Gamma/3G_a$ (μm) |
|---|---|---|---|
| **0.5%** | 1.7 | 1.9 | $3.7\times10^2$ |
| **2%** | $6.5\times10^1$ | $1.9\times10^1$ | $9.8\times10^1$ |

**Table S4. Bacterial strains used in this study.**

| Strain | Genotype | Source |
|---|---|---|
| *V. cholerae* | | |
| JY451 | *vpvC*$^{W240R}$ Δ*vc1807*::*Ptac-mNeonGreen*-Spec$^{R}$ | This study |
| JY452 | *vpvC*$^{W240R}$ Δ*rbmA* Δ*vc1807*::*Ptac-mNeonGreen*-Spec$^{R}$ | This study |
| JY458 | *vpvC*$^{W240R}$ Δ*bap1*Δ*rbmC* Δ*vc1807*::*Ptac-mNeonGreen*-Spec$^{R}$ | This study |
| JY462 | *vpvC*$^{W240R}$ Δ*rbmA*Δ*bap1*Δ*rbmC* Δ*vc1807*::*Ptac-mNeonGreen*-Spec$^{R}$ | This study |
| JY466 | *vpvC*$^{W240R}$ Δ*vpsL* Δ*vc1807*::*Ptac-mNeonGreen*-Spec$^{R}$ | This study |
| JY283 | *vpvC*$^{W240R}$ Δ*pomA* | (8) |
| JY284 | *vpvC*$^{W240R}$ Δ*rbmA* Δ*pomA* | (8) |
| JY285 | *vpvC*$^{W240R}$ Δ*bap1*Δ*rbmC* Δ*pomA* | (8) |
| JY286 | *vpvC*$^{W240R}$ Δ*rbmA*Δ*bap1*Δ*rbmC* Δ*pomA* | (8) |
| JY287 | *vpvC*$^{W240R}$ Δ*vpsL* Δ*pomA* | (8) |
| JY046 | *vpvC*$^{W240R}$ Δ*rbmA* | (1) |
| *S. aureus* | | |
| RN6390b | Standard agr group I prototype | (4) |

### Materials and Methods

**Deconvolution and image processing**. To perform single-cell analysis, we first used Huygens (SVI) to deconvolve the raw image with the measured PSF. Next, we binarized the 3D image layer by layer using the Otsu method (9) and obtain a mask that removed background noise. This step generated many clusters, each containing multiple cells. The key difficulty in segmentation lies in the inferior resolution in the *z*-direction in confocal images as well as imperfect rejection of out-of-focus light, which makes the use of a global threshold inadequate. Therefore, we developed an adaptive local thresholding algorithm for single-cell biofilm image analysis. Specifically, for each cluster, we gradually increased the intensity threshold locally and therefore delete voxels whose intensities are below the threshold. After each local binarization step, we checked whether new clusters are generated. For each newly generated cluster, if its volume was below a defined threshold, we extracted this cluster from the image and considered this cluster as the core of one individual cell. Information on this individual cell was stored and no further action would be performed on this cell until the image-restoration step. If the volume of the cluster was still above the threshold, we performed one of the two following operations depending on the two potential scenarios: 1) This cluster could still be an agglomeration of several cells. The adaptive local thresholding algorithm would be performed on this cluster until it only contained one cell. 2) This cluster represented the core of a single cell that was still connected with neighboring voxels. This cluster would continue shrinking without generating new clusters until it is recognized by the algorithm as a single cell. After the adaptive local thresholding algorithm, we obtained the cores of all individual cells. However, these cores were smaller than the original cells; they did not represent accurately the location, orientation, and size of the original cells. Therefore, we added a volume restoration step in which the deleted voxels in the segmentation step were merged with the nearest core. By performing the virtual shrinking-expansion step, we maintained the accuracy in both segmentation and in measuring the shape of each bacterium.

**Biofilm preparation for rheological measurements.** *V. cholerae* strains were streaked onto LB plates containing 1.5% agar and grown at 37°C overnight. Individual colonies were inoculated into 3 mL of LB liquid medium containing glass beads, and the cultures were grown with shaking at 37°C to mid-exponential phase (5–6 h). Subsequently, the cultures were vigorously vortexed, $OD_{600}$ was measured, and the cultures were back diluted to an $OD_{600}$ of 0.5. 1 μL of this inoculum was spotted onto prewarmed agar plates solidified with 1% agar with M9 medium. Plates were incubated at 37 °C for two days. Sixteen colonies were grown per plate. Biofilm colonies from more than ten plates were agglomerated for each rheological measurement.

**Shear rheology of biofilms.** All rheological measurements were performed with a stress-controlled shear rheometer (Anton Paar Physica MCR502WESP) at 30˚C. For each measurement, 200–400 biofilms were collected with a pipette tip or a razor blade and transferred onto the lower plate of the rheometer (8). After sandwiching the biofilms between the upper and lower plates with a gap size of 0.5 mm, silicone oil (5 cSt at 25˚C, Sigma Aldrich) was applied to enclose the biofilm. Sandblasted surfaces were used for both the upper and lower plates to avoid slippage at the boundary. Oscillatory shear tests were performed with increasing amplitudes of the oscillatory strain $\varepsilon$ from 0.01 to 1500% at a fixed frequency of 1 Hz. The storage modulus $G'$ was extracted with the RheoPlus software as a function of $\varepsilon$. To extract the plateau shear moduli of biofilms, segmented linear fittings were applied to $G'$- $\varepsilon$ curves on a log-log scale. $G'$ varies minimally in the plateau region. We used the fitted $G'$ value at $\varepsilon$ = 1% as the modulus of the biofilm $G_{\mathrm{b}}$ (8, 10). All rheological properties of the biofilm remained roughly constant for at least 48 hours. All rheological measurements were performed in strains lacking motility (Δ*pomA*) to mimic the behavior of cells in the embedded biofilm.

**Shear rheology of agarose gels.** M9 media containing agarose with different concentrations were freshly prepared in 100 mL bottles. The semi-solid medium was heated in a microwave, cooled to ~ 50˚C, and 1 mL of this hot medium was added onto the lower plate of the rheometer preheated to 50˚C. The heated agarose solution was subsequently sandwiched between the two rheometer plates with a gap size of 0.5 mm and sealed with silicone oil. The preparation was cooled to 22˚C using a cooling rate of 2°C/min. Subsequently, the agarose gel was kept at 30°C for measurement. This procedure mimics the sequence of events that agarose gels were exposed to during sample preparation and biofilm growth. Sandblasted parallel surfaces were used with a controlled constant normal force equal to zero during cooling. Oscillatory shear tests were performed in the linear elastic region at a fixed frequency of 1 Hz. 10–20 points in the plateau region of the $G'(\varepsilon)$ curve were averaged to give $G_{\mathrm{a}}$.

**Embedding fluorescent nanoparticles in agarose gels.** To disprove the alternative scenario of crack formation leading to oblate biofilm shapes (11), we performed hydrogel labeling experiments to monitor biofilm growth and potential crack formation in the gel. Appropriate amounts of *V. cholerae* cells constitutively expressing mNeonGreen and 40 nm (diameter) polystyrene nanoparticles (Life Tech. ex/em = 580/605) were mixed in 2% agarose gel so that the final concentrations of cells and nanoparticles were $OD_{600}$ = 0.0002 and $1.4\times10^{12}$ beads/mL, respectively. 50 µL of this mixture was added to the 96-well plate and allowed to solidify for 10 minutes at room temperature.

**Fracture energy measurement of agarose gels.** The oblate biofilm shape is reminiscent of macroscopic penny-shaped cracks commonly observed in brittle materials (12). However, hydrogel labeling experiments show the absence of crack (Fig. S6). Furthermore, we use a scaling analysis (13) here to prove that the embedded biofilms in this study are too small to generate macroscopic cracks (Table S3). To determine the elasto-adhesive length of agarose gel, we performed a common single-edge-notch test based on linear elastic fracture mechanics (14). A pre-notched strip of agarose gel was prepared, in which the pre-notched crack length $c$ is much smaller than the sample width. This agarose strip was clamped with sandblasted paper to the solid rectangular fixture (SRF) feature of the rheometer and stretched at a constant velocity of 50 µm/s. During stretching, the stress $\sigma$ was measured as a function of strain $\varepsilon$. Based on the analytical solution (2), the fracture energy of the gel $\Gamma$ can be calculated as $\Gamma = 4\sigma^2 c/3G_a$, where $G_a$ is the measured shear modulus (13).

**Agarose dissolving experiment.** Following our biofilm growth protocol, we grew *V. cholerae* biofilms inside 2% agarose gels at 30˚C for 1 day to form mature clusters. Subsequently, M9 medium was replaced with 150 μL of agarose dissolving buffer (ADB, Zymo Research) on top of 50 μL gel to dissolve the confining gel. After heating at 40˚C for 1h, at which agarose gels were completely dissolved, the confined biofilm clusters were released. To observe the change in biofilm morphology, we used Syto9 nucleic acid stain (3.34 μM, LIVE/DEAD® BacLight™ Bacterial Viability Kit) to stain bacterial cells before and after gel dissolution.

**Indentation experiment.** To test the mechanical response of different *V. cholerae* mutants, we first prepared a thin agarose sheet between two coverslips with spacers (thickness less than 200 μm). After the gel solidified, we removed the top coverslip, inoculated the agarose surface with *V. cholerae* strains at $OD_{600}$ = 0.2, and allowed cells to settle down and grow for 3 hours to obtain a high surface coverage in order to obtain statistically meaningful data. Subsequently, to mimic the gel deformation process, we used a glass bead (MP Biomedicals, 4 mm in diameter) to deform the agarose substrate covered with isolated bacterial cells and clusters in random orientations. A 60× water objective was used to observe the orientation of both individual cells in isolation and in clusters after indentation. Note that due to the rigidity of agarose gel, the agarose surface formed a much larger deformed curvature than the radius of the spherical indenter. The imaged areas were not in physical contact with the indenter.

**Antibiotic killing assay.** Biofilms of *V. cholerae* strains constitutively expressing mNeonGreen with an initial $OD_{600}$ = 0.0002 were prepared embedded in 2% or 0.5% agarose gels in wells of 96-well plates. After 16h of biofilm growth at 30˚C, the M9 medium on top of the gel was replaced with fresh M9 medium with or without 100 µg/mL tetracycline. After the addition of antibiotics, the embedded biofilms were incubated for another 3h at 30˚C. Subsequently, 20µM of propidium iodide (PI, LIVE/DEAD® BacLight™ Bacterial Viability Kit) was added to stain dead bacterial cells.

**Definition and analysis of order parameters.** After 3D segmentation, based on the coordinates of all constituent cells we determined the major axis of a biofilm using principal component analysis (PCA); subsequent analysis was performed using this internal coordinate system. We defined the shortest axis of the oblate ellipsoid as the *z* axis and recorded the center coordinate (*x*,*y*,*z*) and the unit orientation director $\vec{n}_i$ of each cell. The angle $\theta_i$ with respect to the *z* axis of each cell was defined from the vertical component of $\vec{n}_i$ extracted by $\cos(\theta) = |\vec{n}_i \cdot \vec{z}|$, in which $\vec{z}$ is the unit vector pointing in the *z* direction. To quantify cell ordering, we defined the shell order parameter $S_s$ and bipolar order parameter $S_b$ with respect to the biofilm-gel interface (Fig. S12) for each cell. We used the coordinates of the outmost cells to reconstruct the biofilm-gel interface as a connected web of virtual boundary points. For each boundary point, local curvature $\kappa^{-1}$ and local surface normal $\overrightarrow{n_{\iota,norm}}$ were defined. Each cell was then paired with the nearest boundary point. We defined the local shell order parameter for each cell based on the surface normal direction $\overrightarrow{n_{\iota,norm}}$ and cell direction $\vec{n}_i$ as $S_s = 1 - 3|\vec{n}_i \cdot \overrightarrow{n_{\iota,norm}}|$. The definition of $S_s$ differs slightly from conventional order parameters (15) but it ensures $S_\mathrm{s}$ = 1 for a fully ordered phase and $S_\mathrm{s}$ = 0 for a random phase. Note that $S_\mathrm{s}$ does not contain any information regarding the cell orientation in the plane tangent to the local interface. To characterize the cell orientation in the tangent plane, we used the bipolar order parameter $S_\mathrm{b}$ inspired by the arrangement of LC molecules in a droplet. To define $S_\mathrm{b}$, the center of a cluster was firstly extracted based on the coordinates of all constituent cells. Hence, we defined a new position vector $\vec{r}_i$ for each cell with the center of the coordinate located at the center of the biofilm cluster. The term $S_\mathrm{b}$ was then defined as $S_\mathrm{b} = 1/2(3|\vec{n}_i' \cdot \vec{r}_i'| - 1)$, in which $\vec{n}_i'$ and $\vec{r}_i'$ are the unit vectors of the projection of $\vec{n}_i$ and $\vec{r}_i$ onto the tangent plane. We averaged $S_\mathrm{s}$ and $S_\mathrm{b}$ over cells in the two outmost layers to characterize global cell ordering near the interface, which give $\overline{S_\mathrm{s}}$ and $\overline{S_\mathrm{b}}$, respectively.

***Q*-tensor 2D averaging method.** To determine the average orientation in the biofilm, we calculated the nematic order parameter field $\boldsymbol{Q}$ (16). First, the positions and directions $(x, y, z, n_x, n_y, n_z)$ of each cell were converted to a cylindrical coordinate system $(r, \theta, z, n_r, n_\theta, n_z)$, where $r, z$ axes are defined as shortest axis and longest axis of oblate biofilm in Fig. S12. The bacteria were subsequently binned into sections that were 1 μm ° #1 μm in the $(r, z)$ plane and contained all cells in the $\theta$ direction. In each bin, the $\boldsymbol{Q}$-tensor was calculated, where $\boldsymbol{Q} = < 3\boldsymbol{nn} - \boldsymbol{I} > /2$. Here $\boldsymbol{nn}$ is a dyadic product of the cell direction with itself, and $<\cdot>$ denotes averaging over all cells in a bin. The eigenvalues and eigenvectors of $\boldsymbol{Q}$ in each bin were determined, and the eigenvector corresponding to the largest eigenvalue *S* was taken as the average direction for the cells in that bin. In Fig. S15, we show the eigenvectors projected in the

$(r, z)$ plane, as there is generally little alignment in the $\theta$ direction.

**Continuum modeling approach.** We adopt an energy-based approach to model the evolution of biofilm morphology as it grows. Biofilms lack an inherent tendency to grow in any particular direction or shape. Nonetheless, the introduction of mechanical confinement introduces an energetic penalty on the growth of the biofilm. We hypothesize that the biofilm responds to mechanical confinement by growing in a way that minimizes the total elastic energy of the system. In the initial stages of growth, the confining gel undergoes small to moderate strains and the deformations are purely elastic (denoted as Stage I). As the biofilm continues to grow, large deformation of the agarose gel induces damage (this transition is denoted as Stage II). We show that the observed shape evolution, which leads to non-monotonicity in the biofilms' aspect ratio with respect to the volumetric expansion, can be explained by the onset of damage in the gel.

The modeling approach employed for the different stages is schematically illustrated in Fig. S7, where the transformation of the biofilm into its grown stress-free shape is represented by $\mathbf{F}_g$, and the elastic deformation is represented by $\mathbf{F}_e$. In Stage I, we consider the biofilm as a solid inclusion embedded inside an infinite medium (agarose gel). We assume that the biofilm and the corresponding void in the gel have their stress-free shape (the initial stress-free shape is the same for both) at $\Delta V/V_0 = 0$. As the biofilm increases in volume, at every given $\Delta V/V_0$, it has the freedom to choose its stress-free shape, while the void retains its original stress-free shape. The biofilm chooses its stress-free shape in a way that minimizes the total elastic energy of the system. By carrying out this analysis successively at every $\Delta V/V_0$ we obtain the optimal morphology of the embedded biofilm as a function of the expansion ratio. The results, as discussed in the main text, indicate that the biofilm shape evolution in Stage I is insensitive to the stiffness contrast between the biofilm and the gel ($G_\mathrm{b}/G_\mathrm{a}$), but it does depend on the initial aspect ratio of the biofilm and void (at $\Delta V/V_0 = 0$).

To model Stage II, we introduce a novel and simplified way to account for damage in the gel. In contrast to Stage I, where the stress-free shape of the void in the gel was not allowed to change, in Stage II we allow the void to vary its *effective shape* as a result of damage in the material immediately surrounding the biofilm. With both the void and the biofilm able to change their shapes in Stage II, we reason that their stress-free shapes change to minimize any mismatches in shape, and therefore their stress-free shapes are assumed to be the same. With these assumptions, we search over the space of stress-free shapes to find the one that minimizes the total energy in the system. In contrast to Stage I, biofilm shape evolution in stage II is sensitive to $G_\mathrm{b}/G_\mathrm{a}$. Combining the results of Stage I and Stage II, the model captures the essential elements of the experimental observations. During Stage I the morphology of the biofilm evolves to become more spherical, irrespective of the stiffness contrast. Once the damage initiates, the biofilm-gel system transitions to a different shape evolution pathway depending on the stiffness contrast. If the biofilm is softer than the gel, the model predicts a transition to a higher aspect ratio, and if the biofilm is stiffer than the gel, the biofilm continues to evolve towards the spherical shape.

**Computational implementation.** The mechanics model summarized above was computationally implemented via the commercial nonlinear Finite Element software COMSOL 5.3a. For simplicity, we limited the analysis to 2D by considering plane-strain conditions and focusing only on elliptical stress-free shapes of the biofilm and the void. Both biofilm and gel were modeled as nearly incompressible neo-Hookean solids, whose strain energy density function reads

$$W(\mathbf{F}) = \frac{G}{2}(\mathbf{F}:\mathbf{F} - 3) - G\ln(\det(\mathbf{F})) + \frac{K}{2}(\ln(\det(\mathbf{F})))^2, \tag{S1}$$

where $G$ and $K$ denote the shear modulus and bulk modulus, respectively, and $\mathbf{F} = \mathbf{F}_e\mathbf{F}_g$ is the deformation gradient. The ratio $K/G$=$10^3$ was set to maintain a nearly incompressible response of the material. To eliminate boundary effects, we set the radius of the gel environment as 1000 times the semi-major axis of the elliptical inclusion (Fig. S7A). For computational efficiency, only a quarter of the geometry with symmetric boundaries was modeled; in addition, we employed a highly refined mesh near the biofilm and coarser mesh far away from the biofilm to balance the accuracy and computation efficiency. Mesh sensitivity analysis confirms that simulation results are mesh-size independent.

Considering the deformation illustrated in Fig. S7, we can express the growth tensor as

$$\mathbf{F}_g = \begin{bmatrix} \alpha & 0 & 0 \\ 0 & \beta & 0 \\ 0 & 0 & 1 \end{bmatrix},$$

the volume growth ratio as $\Delta V/V_0 = \alpha\beta - 1$, and the stress-free aspect ratio as $b'/a'$ $= (\beta b)/(\alpha a)$. By choosing $\mathbf{F}_g$ we computationally prescribe the stress-free shape of the biofilm at a given volume expansion.

In Stage I (purely elastic deformation), for every $\Delta V/V_0$ we computationally scan over the space of growth tensors $\mathbf{F}_g$ (adhering to the constraint on $\Delta V/V_0$) for the biofilm, while the void retains its stress-free shape, and calculate the total energy of the system for each case. Following the model outlined in the previous section, we then select the growth tensor that minimizes the total energy (example case is shown in Fig. S8A). We repeat this two-step procedure successively for different volume expansions ($\Delta V/V_0$) to obtain the morphological evolution of the biofilm with increasing volume (Fig. S8B).

In Stage II (damage transition), for every expansion ($\Delta V/V_0$) we computationally scan over the space of growth tensors $\mathbf{F}_g$ (adhering to the constraint on $\Delta V/V_0$) for the biofilm, while requiring that, through damage, the void also assumes the same stress-free shape as the biofilm (example case is shown in Fig. S8C). Following the same two-step computational procedure as outlined in Stage I, we obtained the curve as shown in Fig. 8D. Whenever the damage transition takes place, the biofilm-gel morphology is expected to jump to the corresponding location on the Stage II curve for the same $\Delta V/V_0$. Sensitivity of Stage II to the stiffness ratio is shown in Fig. S9.

**Supplementary Movies S1-4**

**Movies S1| *z*-scan of *V. cholerae* biofilms from bottom to top.** *z*-Scan of two 15h-old Δ*rbmA V. cholerae* biofilms grown in 2% agarose gel (from bottom to top), one aligning approximately perpendicular and the other one parallel to the imaging plane. The strain constitutively expresses mNeonGreen. The field of views are 56 μm × 40 μm and 56 μm × 45 μm, respectively.

**Movies S2| 3D view and reconstruction of a biofilm cluster.** Three-dimensional view and single-cell reconstruction of a mature Δ*rbmA* biofilm cluster. Colors denote cell orientation, which is defined as the angle between the cell axis and the *z* axis. The field of each view is 60 μm × 52 μm.

**Movies S3| Growth of *V. cholerae* biofilms in 2% and 0.3% gels.** A time-lapsed cross-sectional view of a Δ*rbmA V. cholerae* biofilm growing inside 2% and 0.3% agarose gel, respectively. Imaging began 2 h after seeding bacterial cells in the agarose gel, and the total duration of the movie is 16 h. The field of view is 33 μm × 40 μm.

**Movies S4| Growth of a reconstructed biofilm cluster with cell ordering.** Time course of segmented reconstruction of a biofilm cluster growing inside 2% agarose gel with each cell colored by shell order parameter $S_s$ (Top) and bipolar order parameter $S_b$ (Bottom).
